# Cooperative and structural relationships of the trimeric Spike with infectivity and antibody escape of the strains Delta (B.1.617.2) and Omicron (BA.2, BA.5, and BQ.1)

**DOI:** 10.1101/2023.03.06.531335

**Authors:** Anacleto Silva de Souza, Robson Francisco de Souza, Cristiane Rodrigues Guzzo

## Abstract

Herein, we simulated the trimeric Spike of the variants B.1.617.2, BA.2, BA.5 and BQ.1 for 300 ns. We derived mechanisms by which the substitutions K417N, L452R, N444T and N460K may favor resistance to neutralizing antibodies. The K417N and L452R contribute to the expansion of the networks of hydrogen bonding interactions with neighboring residues, decreasing their capacity to interact with neutralizing antibodies. The Spike^BQ.1^ possesses two unique K444T and N460K mutations that expand the network of hydrogen bonding interactions. This lysine also contributes one novel strong saline interaction and both substitutions may favor resistance to neutralizing antibodies. We also investigated how the substitutions D614G, P681R, and P681H impact Spike structural conformations and discuss the impact of these changes to infectivity and lethality. The prevalent D614G substitution plays a key role in the communication between the glycine and the residues of a β-strand located between the NTD and the RBD, impacting the transition between up- and down-RBD states. The P681R mutation, found in the Delta variant, favors intra- and inter-protomer correlations between the subunits S1 and S2. Conversely, in Omicron sub-variants, P681H decreases the intra- and inter-protomer long-range interactions within the trimeric Spike, providing an explanation for the reduced fusogenicity of this variant. Taken together, our results enhance the knowledge on how novel mutations lead to changes in infectivity and reveal mechanisms by which SARS-CoV-2 may evade the immune system.

## Introduction

The rapid evolution of SARS-CoV-2 is dominated by the accumulation of mutations in the viral genome, impacting many aspects of the biology of the virus, such as its speed of propagation, pathogenicity and infectivity (de Souza, de Freitas Amorim, Guardia, Dos Santos, Ulrich, et al., 2022). SARS-CoV-2 variants whose emergence is associated with important waves of infection are often designated as variants of concern (VOCs). In order to allow monitoring of the spread of their genetic variants, VOCs are classified into Nextstrain clades and Pango lineages (GISAID, 2022a). To facilitate discussions for both technical and non-scientific audiences, the World Health Organization (WHO) recommends the use of letters from the Greek alphabet to match the order of appearance of the SARS-CoV-2 VOCs, such as Alpha, Beta, Gamma, Delta, and Omicron (WHO, 2022b). Thus, VOCs have been classified as Alpha (pango lineage, B.1.1.7; nextstrain clade, 20I), Beta (pango lineage, B.1.351; nextstrain clade, 20H), Gamma (pango lineage, P.1; nextstrain clade, 20J), Delta (pango lineage, B.1.617.2; nextstrain clade, 21A, 21I, 21J), and the globally predominant strain, the Omicron variant (pango lineage, B.1.1.529; nextstrain clade 21M). The Omicron VOCs include several sub-lineages such as BA.1 (nextstrain clade 21M), BA.2 (nextstrain clade 21L), BA.4 (nextstrain clade 22A), BA.5 (nextstrain clade 22B), BA.2.12.1 (nextstrain clade 22C), BA.2.75 (nextstrain clade 22D), BQ.1 (nextstrain clade 22E), and XBB (recombination between BA.2.10.1 and BA. 2.75, nextstrain clade 22F) (GISAID, 2022a). Although the Delta variant has a high degree of transmissibility and infectivity, the Omicron variant quickly became predominant worldwide, thus proving to be even more transmissible than Delta (de Souza, de Freitas Amorim, Guardia, Dos Santos, Ulrich, et al., 2022). BA.1 and BA.2 are less lethal in vaccinated individuals (WHO, 2022a), but are able to escape the immune system (Gobeil et al., 2022; Stalls et al., 2022). BA.4, BA.5, and BQ.1 are currently the most predominant strains worldwide and they are related to the most recent surges in cases of COVID-19 (Callaway, 2022; GISAID, 2022b).

When SARS-CoV-2 variants are compared, mutations can be found in any gene, but a higher number of mutations concentrate in the gene region that encodes the Spike protein (de Souza, de Freitas Amorim, Guardia, Dos Santos, Ulrich, et al., 2022). Spike is a homotrimer anchored in the viral membrane composed by S1 and S2 subunits, each performing a different role in the viral infection (de Souza, de Freitas Amorim, Guardia, Dos Santos, Dos Santos, et al., 2022; de Souza, de Freitas Amorim, Guardia, Dos Santos, Ulrich, et al., 2022). The S1 subunit contains the N-terminal domain (NTD), the receptor binding domain (RBD) that play key roles in recognition of the hACE2, SD1 and SD2 domains (de Souza, de Freitas Amorim, Guardia, Dos Santos, Dos Santos, et al., 2022; de Souza, de Freitas Amorim, Guardia, Dos Santos, Ulrich, et al., 2022; Sironi et al., 2020; Souza et al., 2020). The RBD may be present in two conformational states, up or down. In its up-conformational state, the RBD is capable of binding to hACE2. Conversely, the S2 subunit mediates the fusion between virus and host membranes (de Souza, de Freitas Amorim, Guardia, Dos Santos, Ulrich, et al., 2022; Zhu et al., 2021). Spike may be recognized and cleaved by the furin protease in the junction connecting S1 with S2 (S1/S2 site), at position R682-R685 (RRAR_685_↓) (**Figures S1 and S2**) (de Souza, de Freitas Amorim, Guardia, Dos Santos, Ulrich, et al., 2022; Wu & Zhao, 2020; Y. Zhang et al., 2022). In addition, transmembrane Serine Protease 2 (TMPRSS2) and Cathepsin isoforms also recognize and cleave the S2’ site (KPSKR_815_↓) (Bosch et al., 2008; de Souza, de Freitas Amorim, Guardia, Dos Santos, Dos Santos, et al., 2022; Essalmani et al., 2022; Hoffmann et al., 2020; Zang et al., 2020; M.-M. Zhao et al., 2022). After hACE2 interaction and S2’ cleavage, the S2 subunit undergoes a major structural change to expose the fusion loop, which mediates the fusion of virus envelope and host cell membrane, allowing the formation of a pore for the release of viral RNA into the cell cytoplasm (Jackson et al., 2022).

Our previous studies showed that the Spike^VOCs^ have similar binding affinities to hACE2, but these are higher than that of Spike^WT^ (de Souza, de Freitas Amorim, Guardia, Dos Santos, Dos Santos, et al., 2022; de Souza, de Freitas Amorim, Guardia, Dos Santos, Ulrich, et al., 2022). Previously results have shown that substitutions K417N, S477N, T478K, E484K and N501Y cause an increase in binding affinity between RBD and hACE2 (Chen et al., 2021; de Souza, de Freitas Amorim, Guardia, Dos Santos, Dos Santos, et al., 2022; de Souza, de Freitas Amorim, Guardia, Dos Santos, Ulrich, et al., 2022; de Souza et al., 2021; Tian et al., 2021). The substitutions L452R and F486V found in BA.4 and BA.5 play a key role in the decrease of neutralizing antibody recognition (Cao, Yisimayi, et al., 2022; Tuekprakhon et al., 2022).

Previous experimental studies have shown that, in addition to high binding affinity, other biochemical variables are related to the enhanced viral infection and increased transmissibility of VOCs (de Souza, de Freitas Amorim, Guardia, Dos Santos, Dos Santos, et al., 2022; de Souza, de Freitas Amorim, Guardia, Dos Santos, Ulrich, et al., 2022), including: **(1)** increased structural flexibility that would make it difficult to be recognized by neutralizing antibodies in Spike^BA.1^ (de Souza, Amorim, et al., 2022; Stalls et al., 2022); **(2)** independence of TMPRSS2 cleavage, leading to changes in cell tropism (de Souza, Amorim, et al., 2022; Meng et al., 2022; Willett et al., 2022); **(3)** increased spike density on the viral surface particle (de Souza, de Freitas Amorim, Guardia, Dos Santos, Ulrich, et al., 2022; Yurkovetskiy et al., 2020; L. Zhang et al., 2020); **(4)** improved exposure of RBD to interact with hACE2 (Benton et al., 2021; de Souza, Amorim, et al., 2022; Mansbach et al., 2021a; Teruel et al., 2021). Although the Omicron variant is more transmissible than the Delta variant, the evolution of the mortality rates for these variants suggests that the Omicron variant is less lethal than the Delta variant (Rahmani & Rezaei, 2022; C. Wang et al., 2022).

In view of the emergency of the BA.5 and BQ.1 variants and the replacement of the B.1.617.2 and BA.2 by these new variants (GISAID, 2022b), we investigate how mutations are related to conformational differences among Spike^B.1.617.2^, Spike^BA.2^, Spike^BA.5^ and Spike^BQ.1^. We performed molecular dynamics simulations of the trimeric ectodomains of these Spike variants for 300 ns. We analyzed the impact of substitutions that mediate resistance to neutralizing antibodies by calculating hydrogen bonding occupancies. To assess the impact of mutations on the structural conformations of the Spikes, we calculated the values of root-mean-square deviation (RMSD) and root-mean-square fluctuation (RMSF) and performed a principal component analysis (PCA) of the structural ensembles. These analyzes allowed us to map important variations in conformational flexibilities, especially in the NTD and the RBD. We also obtained the dynamical cross-correlation matrix (DCCM) for the simulated trajectories, which showed that changes to inter- and intra-protomer communication may be responsible for reduced efficiency of the fusogenic process. Since SARS-CoV-2 VOCs may be more pathogenic, transmissible or more able to evade the immune system, detailed understanding of how the mutations of the Spike protein impact its structure and function is essential to guide prediction of fitness gains and enable rational development of updated vaccines in COVID-19 therapy.

## Materials and Methods

### Homology modeling

In order to investigate how molecular dynamics may influence transmissibility and infectivity, we simulated the trimeric Spike^B.1.617.2^, Spike^BA.2^, Spike^BA.5^ and Spike^BQ.1^ ectodomains in a water-filled box. The available structures have mutations and/or unresolved regions. In order to correct these problems, we used cryogenic electron microscopy (Cryo-EM) of the Spike^BA.1^ structure (PDB ID 7WK2) as a template to obtain the corrected structures in the Swiss-Model web server (Waterhouse et al., 2018). Initially, the genomic sequences of the B.1.617.2 (GISAID, accession code EPI_ISL_15728122), BA.2 (GISAID, accession code EPI_ISL_12905607), BA.5 (GISAID, accession code EPI_ISL_14439279) and BQ.1 (GISAID, accession code EPI_ISL_15731012) were translated in Translate tools (available on the Expasy web server) to obtain the primary sequence of the corresponding Spikes proteins. All primary sequences were submitted in the Swiss Model web service (Bertoni et al., 2017; Bienert et al., 2016; Waterhouse et al., 2018), ignoring the transmembrane regions. Furthermore, we considered the unglycosylated state of Spike because the difference between glycosylated and unglycosylated Spike is smaller than seen between different Spike variants (Zimmerman et al., 2021). In order to compare all molecular dynamics simulations, we used only the structures in the down conformation (all 3D models are available in **supplementary file**).

### Setup for molecular dynamics simulations

In order to study the structural features of the Spikes, initially we determined the protonation states of ionizable residues in an implicitly aqueous environment at pH 7.0 using the PROPKA module implemented in the Maestro program (academic version v. 2021-4, by Schrödinger) (Schrodinger, 2021). Thus, **(1)** all glutamic and aspartic residues were represented as non-protonated; **(2)** arginine and lysine residues were assumed to have a positive charge; **(3)** in the N- and C-terminal regions, the amino and carboxyl groups have been converted to charged groups; **(4)** the histidines were protonated according to prediction of PROPKA and manual inspection, such as: **Spike^B.1.617.2^**, δ-tautomer (H75, H664, H1067, H1073, H1092, H1097, and H1110) and ε-tatomer (H58, H78, H216, H254, H528, H634, H1057) and charged histidine in H155 residue; **Spike^BA.2^**, δ-tautomer (H72, H251, H511, H960, H1054, H1064, H1070, 1089, 1094, 1107) and ε-tatomer (H55, H75, H152, H213, H525, H631, H687); **Spike^BA.5^**, δ-tautomer (H72, H509, H629, H958, H1052, H1062, H1068, H1087, H1092, H1105) and ε-tatomer (H55, H150, H211, H249, H523, H685); **Spike^BQ.1^**, δ-tautomer (H72, H149, H508, H628, H957, H1051, H1061, H1067, H1086, H1091, H1104) and ε-tatomer (H55, H210, H248, H522, H684). In **Spike^B.1.617.2^**, the disulfide bridges were considered between residue pairs: C140/C175, C300/C310, C345/C370, C388/C441, C400/C534, C489/C497, C547/C599, C626/C658, C671/C680, C747/C769, C752/C758, C1041/C1052, C1091/C1135. In **Spike^BA.2^**, the disulfide bridges were considered between residue pairs: C137/C172, C297/C307, C342/C367, C385/C438, C397/C531, C486/C494, C544/C596, C623/C655, C668/C677, C744/C766, C749/C755, C1038/C1049, C1088/C1132. In **Spike^BA.5^**, the disulfide bridges were considered between residue pairs: C135/C170, C295/C305, C340/C365, C383/C436, C395/C529, C484/C492, C542/C594, C621/C653, C666/C675, C742/C764, C747/C753, C1036/C1047, C1086/C1130. In **Spike^BQ.1^**, the disulfide bridges were considered between residue pairs: C135/C169, C294/C304, C339/C364, C382/C435, C394/C528, C483/C491, C541/C593, C620/C652, C665/C674, C741/C763, C746/C752, C1035/C1046, C1085/C1129. All structures were previously minimized in Chimera tools, using steepest descent steps of 500, steepest descent step size of 0.02 Å, conjugate gradient steps of 10 and update interval of 10.

All molecular dynamics simulations were performed using GROMACS, v. 2022.1 (Abraham et al., 2015; Lindahl et al., 2001; Pronk et al., 2013; Van Der Spoel et al., 2005), using the OPLS-AA force field (Robertson et al., 2015). All systems were then explicitly solvated with TIP3P water models in a triclinic box and neutralized, maintaining the concentration of 150 mM NaCl and minimized until reaching a maximum force of 10.0 kJ/mol or a maximum number of steps in 50,000. The systems were consecutively equilibrated in isothermal-isochoric (NVT) ensembles by 2 ns (number of steps and intervals of 1.000,000 and 2 fs, respectively) and isothermal-isobaric (1 bar, 310 K, NpT) by 2,000 ps (number of steps and intervals of 1,000,000 and 2 fs, respectively). All simulations were then performed in a periodic triclinic box considering the minimum distance of 1.2 nm between any protein atom and the walls of the box. Molecular dynamics runs were performed in isothermal-isobaric (1 bar, 310 K, NpT) by 300 ns (number of steps and intervals of 150,000,000 and 2 fs, respectively) in the NpT ensemble. We use the leap-frog algorithm to integrate Newton equations. We selected LINCS (LINEar Constraint Solver) that satisfies the holonomic constraints, whose number of iterations and order were 1 and 4, respectively. We use neighbor search grid cells (Verlet clipping scheme, frequency to update 20-step neighbor list, and clipping distance for 12 Å^2^ short-range neighbor list). In the van der Waals parameters, we smoothly shift the forces to zero between 10 and 12 Å. In electrostatic Coulomb, we use Particle-Mesh Ewald (PME) fast and smooth electrostatics for long-range electrostatics. In addition, we set the distance for the Coulomb cut to 12 Å, order of interpolation for PME to value 4 (cubic interpolation) and grid spacing for Fast Fourier Transform (FFT) to 1.6 Å. In temperature coupling, we use velocity rescheduling with a stochastic term (V-resale, modified Berendsen thermostat). After obtaining two coupling groups (protein and water/ions), we consider the time constant (0.1 ps) and 310 K as the reference temperature. In the pressure coupling (NpT ensembles), we use the Parrinello-Rahman barostat (isotropic type, time constant of 2 ps, reference pressure of 1 bar and isothermal compressibility of 4.5×10^-5^ bar^-1^). We saved the compressed coordinates every 10.0 ps. In molecular dynamics calculations, we use periodic boundary conditions in xyz coordinates (3D space). We then calculated the percent hydrogen bond occupancy (all frames, considering cut-off of distance and angle of 4 Å and 20°, respectively), root-mean-square deviation (RMSD) and root-mean-square fluctuation (RMSF) using the GROMACS modules and Visual Molecular Dynamics (VMD) (Humphrey et al., 1996). Analysis of C_α_ dynamical cross-correlation matrix (DCCM) was performed using the R-based package Bio3d (Grant et al., 2006).

### Principal component analysis

Principal component analysis was calculated using the GROMACS modules, g_covar and g_anaeig. Using simulated molecular dynamics trajectories, we determined the average position of the Spike backbone atoms and calculated covariance matrix fitting non-mass weighted. Covariance matrices were constructed using the backbone of the Spike^B.1.617.2^ (30,591×30,591), Spike^BA.2^ (30,537×30,537), Spike^BA.5^ (30,483×30,483), and Spike^BQ.1^ (30,456×30,456). From covariance matrices, we diagonalized them to obtain the eigenvalues (whose sum of all eigenvalues were of 510.3, 572.0, 561.8, and 385.8 nm²) and, afterwards, orthogonal eigenvectors. Using the first eigenvalue, we determined the eigenvector 1 (named principal component 1, PC1) that represents the first main movement detected along molecular dynamics trajectories. We projected molecular dynamics trajectories of the Spike^B.1.617.2^, Spike^BA.2^, Spike^BA.5^, and Spike^BQ.1^ in PC1, skipping 300 frames, to obtain 100 snapshots and to visualize the structural changes of these proteins in VMD software (**supplementary files S1-S4**). In order to quantify the conformational changes of the main movements of the Spike in 310 K, we calculated the entropy by Schlitter method (Schlitter, 1993) (ΔS^Schlitter^), using g_anaeig of the GROMACS.

## Results and Discussion

### Sequence and structural variations among Spike^VOCs^

Several mutations in the Spike^VOCs^ are related to increased binding affinity, increased infectivity, immune system escape and increased transmissibility of SARS-CoV-2 (de Souza, de Freitas Amorim, Guardia, Dos Santos, Ulrich, et al., 2022). Multiple sequence alignment of Spike^WT^, Spike^B.1.617.2^, Spike^BA.2^, Spike^BA.5^, and Spike^BQ.1^ proteins (**Figure S1** and **S2**) shows that Spike^B.1.617.2^ presents mutations in NTD (T20R, G75R, D80Y, T95I, G142D), RBD (L452R, T478K), in the SD1 subdomain (T547K, D614G, P681R), the SD2 subdomain (D614G, H655Y), and in S2 subunit (D950N and V1264L). Conversely, Spikes of BA.2, BA.5 and BQ.1 share the following mutations when compared to Spike^WT^: NTD (T20I, del25/27, A28S, G339D), RBD (S371F, S373P, S375F, T376A, D405N, R408S, K417N, S477N, T478K, E484A, Q498R, N501Y, Y505H), SD1 subdomain (D614G, H655Y, N679K, P681H, N764K, D796Y). In Spike^BA.2^, residues H69, V70, N440, L452, F486 are the same as the residues in the wild type, but there is the substitution Q493R only found in this Omicron sub-variant. Spike^BA.5^ and Spike^BQ.1^ have the L452R substitution and deletions of the residues H69 and V70. Comparing the sequences of Spike^BQ.1^ and Spike^BA.5^, we observed few differences such as a deletion at Y145 in BQ.1 and the substitutions K444T and N460K. These three mutations are only found in Spike^BQ.1^.

In order to investigate how mutations are related to structural changes of different Spike of VOCs, we compared the backbone root-mean-square deviation (RMSD) of the protomers of Spike^B.1.617.2^ (from Q14 to S1146), Spike^BA.2^ (from Q14 to S1144), Spike^BA.5^ (from Q14 to S1142) and Spike^BQ.1^ (from Q14 to S1141) (**Figure 1**). The average RMSD values of Spike^B.1.617.2^, Spike^BA.2^, Spike^BA.5^ and Spike^BQ.1^ converged with ∼5.0, ∼5.1, ∼5.0 and ∼4.9 Å, respectively. Our previous study resulted in RMSD values of ∼5 Å for Spike^BA.1^ and ∼4 Å for Spike^WT^ (de Souza, Amorim, et al., 2022). Therefore, Spike^B.1.617.2^, Spike^BA.2^, Spike^BA.5^, and Spike^BQ.1^ visit a greater number of different structural conformations, similar to Spike^BA.1^, when compared with the wild type.

**Figure 1.**
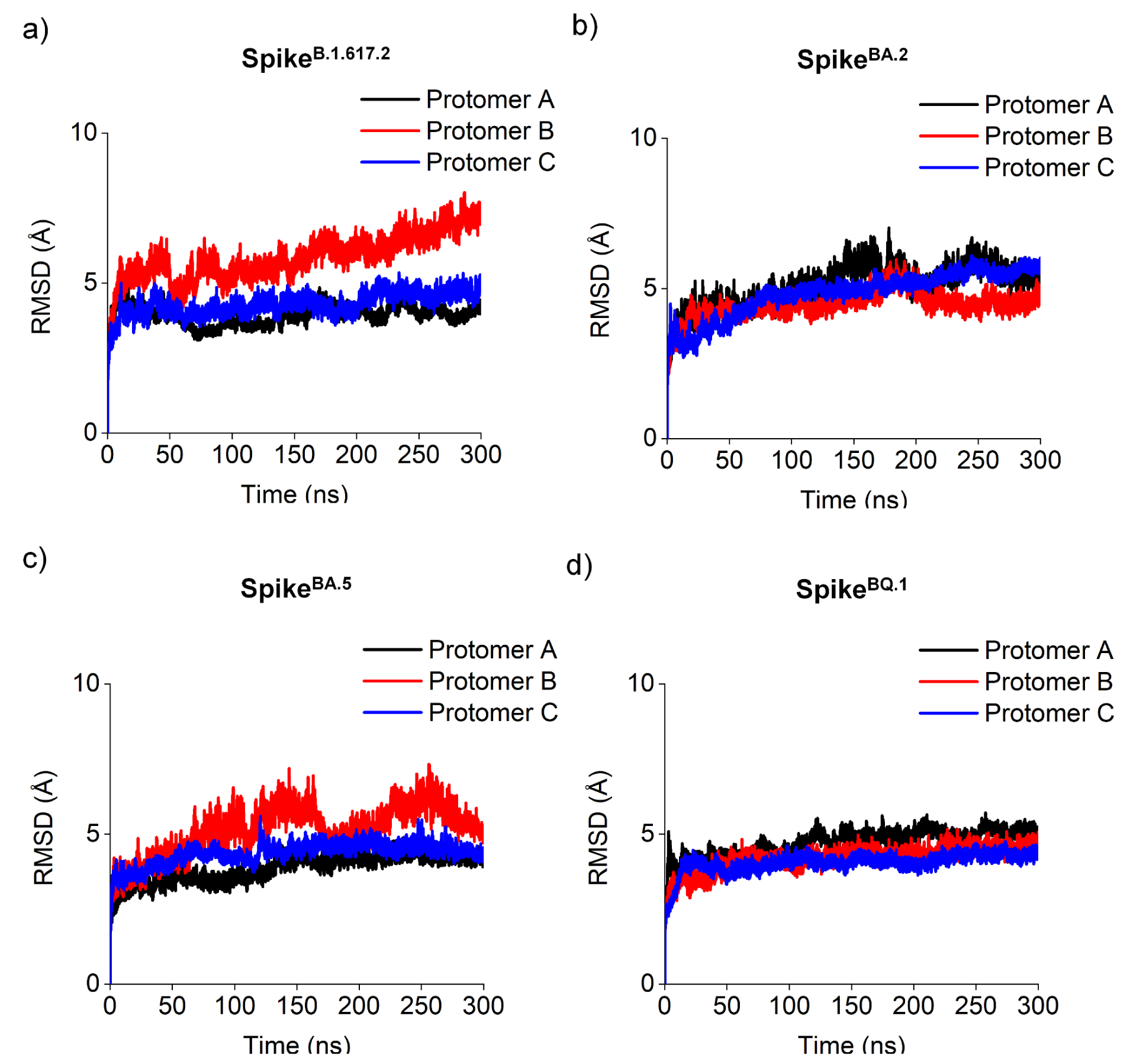
Structural aspects of Spike^B.1.617.2^, Spike^BA.2^, Spike^BA.5^, and Spike^BQ.1^. Backbone root-mean-square deviation (RMSD) calculated from the molecular dynamics (MD) simulations of the **a)** Spike^B.1.617.2^**, b)** Spike^BA.2^, **c)** Spike^BA.5^, and **d)** Spike^BQ.1^.

Next, we calculated the backbone root-mean-square fluctuation (RMSF) of the amino acid residues of Spike^VOCs^ (**Figure 2**). We attributed a RMSF cut-off value of 3 Å and classified the residues with high conformational flexibility as those with an average RMSF greater than 3 Å. To facilitate the location of amino acid residues in the Spike, we have mapped the residue numbering based on the domain architecture of the Spike^WT^ protein, as shown in the **Figure 2a**. Spike^B.1.617.2^ mutations are located in NTD (T20R, G75R, D80Y, T95I, G142D), RBD (L452R, T478K), SD1 (T547K, D614G, P681R), SD2 (D614G, H655Y), on HR1 (D950N) and HR2 (V1264L) (**Figure S1** and **S2**). These mutations influence the structural flexibility of Spike^B.1.617.2^, as shown by the high RMSF value in NTD (14-23, 71-74, 113, 145-155, 164-165, 167, 212-218, 248-257), in RBD (474-488), in SD2 subdomain (622-626), in proximity of the S1/S2 cleavage site (679-690), in proximity of the S2’ cleavage site (810-812), between the FP and HR1 domains (836-846), in HR1 domain (941-942), and in HR2 domain (1140-1146) (**Figure 2b**). The residues with high RMSF values in Omicron sub-variants are: **(1) BA.2**, in NTD (14-20, 66-76, 142-150, 161-163, 210, 243-255), in RBD (468-486), in SD2 subdomain (620-623), in proximity of the S1/S2 site (675-685), in proximity of the S2’ site (806-810), between the FP and HR1 domains (842), in HR1 domain (938, 940-941), and in HR2 domain (1138-1144) (**Figure 2c**); **(2) BA.5**, in NTD (14-21, 67-73, 139-146, 152, 175-180, 207-210, 240-253), in RBD (469-482), in proximity of the S1/S2 site (673-683), in proximity of the S2’ site (803-808), in FP (827), between FP and HR1 domains (830-841), and in HR2 domain (1134-1142) (**Figure 2d**); **(3) BQ.1**, in NTD (14-21, 68-71, 140-145, 177, 206-208, 241-251), in RBD (470-479), in SD2 subdomain (616-620), in proximity of the S1/S2 site (673-680), in proximity of the S2’ site (803-807), and in HR2 domain (1135-1141) (**Figure 2e**). In our previous study, Spike^BA.1^ showed high RMSF values for residues in NTD (14-21, 70-75, 142-149, 175-179, 208-215, 243-255), in RBD (442-444 and 468-485), in SD2 subunit (599-603, 635 and 654), in proximity of the S1/S2 site (672-690), in FP (828-838), in HR1 domain (938-944), and in HR2 domain (1139-1144)(de Souza, Amorim, et al., 2022; Sakkiah et al., 2020). When compared with Spike^BA.1^ (de Souza, Amorim, et al., 2022), our results for Spike^BA.2^, Spike^BA.5^, and Spike^BQ.1^ show that these proteins are less flexible in the S2 subunit, but more flexible in the S1 subunit (mainly in the NTD and RBD). In our previous study, we defined two regions of the Spike RBD interaction interface with human receptor Angiotensin-converting enzyme 2 (hACE2), E1 (residues K417, L455, F456, and 470-490) and E2 (residues 446-453 and 493-505), which have different physicochemical features that impact the interaction with hACE2 (Souza et al., 2020). E1 makes mostly hydrophobic interactions, while E2 interacts mostly with hydrophilic interactions with hACE2 (Souza et al., 2020). The Spikes of BA.2, BA.5 and BQ.1 share the same mutations in NTD (T20I, del25/27, A28S, G339D), in RBD (S371F, S373P, S375F, T376A, D405N, R408S, K417N, S477N, T478K, E484A, Q498R, N501Y, Y505H), and SD1 (D614G, H655Y, N679K, P681H, N764K, D796Y). It is known that substitutions in E1 region (K417N, S477N, T478K, E484K) and E2 region (N501Y) contribute to increase the binding affinity between RBD and hACE2 (Chen et al., 2021; de Souza, de Freitas Amorim, Guardia, Dos Santos, Dos Santos, et al., 2022; de Souza, de Freitas Amorim, Guardia, Dos Santos, Ulrich, et al., 2022; Tian et al., 2021). All results are in total agreement with several previous studies, which show high RMSF values in the residues 470-480 (in E1 region), interacting or not with hACE2 (de Souza, Amorim, et al., 2022; de Souza, de Freitas Amorim, Guardia, Dos Santos, Dos Santos, et al., 2022; Sakkiah et al., 2020; Souza et al., 2020). The L452R and F486V substitutions decrease the sensitivity of neutralizing antibody recognition in BA.4 and BA.5 sub-variants (Cao, Yisimayi, et al., 2022; Tuekprakhon et al., 2022). Overall, we did not observe significant RMSF differences in the S1 subunit of the Spike^BA.1^ (de Souza, Amorim, et al., 2022), Spike^B.1.617.2^, Spike^BA.2^, Spike^BA.5^, and Spike^BQ.1^. However, we noted a significant decrease of the RMSF values in the proximity of the S1/S2 and S2’ cleavage sites of the Spike^BA.5^ and Spike^BQ.1^ when compared with other Spike VOCs studied in this paper (de Souza, Amorim, et al., 2022). As described in more details below, we hypothesize that this decrease in RMSF shows a reduction in mobility of residues. We believe that this effect may be affected by diminished cooperativity induced by mutations shared by the Spike proteins of Omicron sub-variants. This loss of cooperativity between Spike protomers could be responsible for a decrease in fusogenicity.

**Figure 2.**
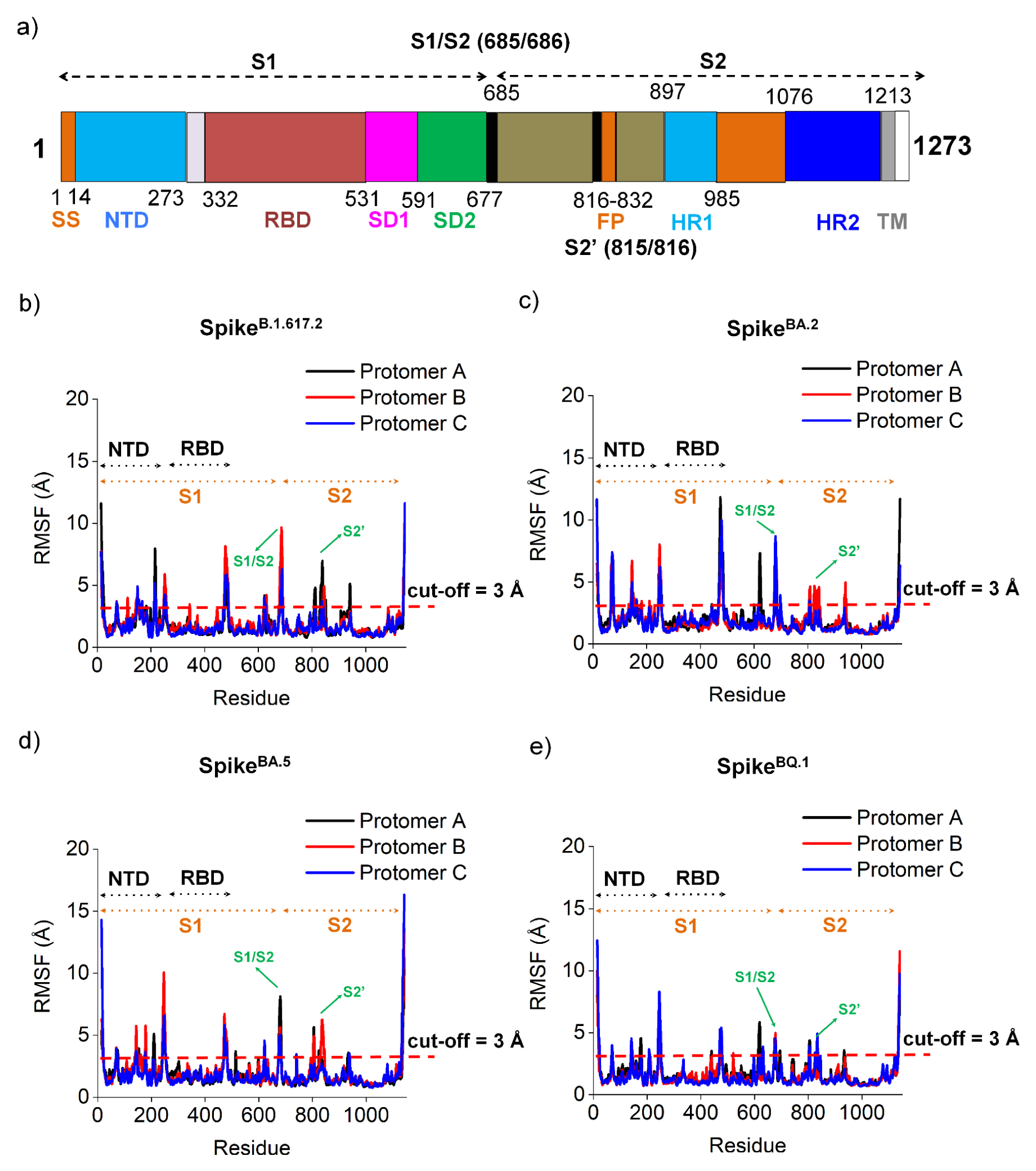
Structural flexibility of the trimeric Spike^B.1.617.2^, Spike^BA.2^, Spike^BA.5^, and Spike^BQ.1^. **a)** Spike^WT^ amino acid sequence showing the S1 and S2 subunits and the cleavage sites S1/S2 and S2’ (de Souza, de Freitas Amorim, Guardia, Dos Santos, Ulrich, et al., 2022). Backbone root-mean-square fluctuation (RMSF) calculated from the molecular dynamics (MD) simulations of the **b)** Spike^B.1.617.2^**, c)** Spike^BA.2^, **d)** Spike^BA.5^, and **e)** Spike^BQ.1^.

In order to identify the most important movements within the Spike trimers, we performed a principal component analysis (PCA) from the diagonalization of the covariance matrix of Spike^B.1.617.2^, Spike^BA.2^, Spike^BA.5^ and Spike^BQ.1^. From the eigenvalues estimated by PCA, we calculated the Schlitter entropy (ΔS^Schlitter^) as an approximation for the global entropy of the conformers of Spike^B.1.617.2^, Spike^BA.2^, Spike^BA.5^, and Spike^BQ.1^, obtaining values of 74.8, 74.6, 75.3, and 74.3 kJ.mol^-1^.K^-1^, respectively. The trimeric Spike^BA.1^ previously reported entropy was 75.1 kJ.mol^-1^.K^-1^ *(de Souza, Amorim, et al., 2022)*. Therefore, the values of entropy for the analyzed variants were similar among variants but significantly higher than the entropy of Spike^WT^ calculated in our previous study (72.5 kJ.mol^-1^.K^-1^) (de Souza, Amorim, et al., 2022). Using the first eigenvector (PC1, first principal component), we verified that main conformational changes of the Spike^B.1.617.2^, Spike^BA.2^, Spike^BA.5^, and Spike^BQ.1^ are localized majorly in NTD and RBD (**Figures 3** and **4**). A broad spectrum of conformations in these regions are important because they may decrease recognition by neutralizing antibodies against SARS-CoV-2 (Yang & Du, 2021). Indeed, the high fluctuation in nearby loops 144-155 and the adjacent loops 246-260 observed in different VOCs causes decreased neutralizing antibody activity (Cai et al., 2021). In general, high structural changes of the NTD and RBD decrease binding affinity to neutralizing antibodies, contributing to the immune system evasion. SARS-CoV-2 VOCs are able to evade the neutralizing antibodies from convalescent or/and immunized patients (Casadevall et al., 2021; de Souza, de Freitas Amorim, Guardia, Dos Santos, Ulrich, et al., 2022; Khan et al., 2021, 2022). Furthermore, deletions 242-244 in Spike cause structural changes in the NTD, also decreasing neutralizing antibody efficiency (Cai et al., 2021). Notably, when compared with the original strain, there are more mutations in NTD and RBD than in the other domains (**Figure S2**). Since NTD and RBD contain important epitopes for neutralizing antibody recognition, mutations in these domains cause a reduction of vaccine-induced neutralizing antibody recognition (Cao, Yisimayi, et al., 2022; Dejnirattisai et al., 2022; de Souza, de Freitas Amorim, Guardia, Dos Santos, Ulrich, et al., 2022; Gobeil et al., 2022; Stalls et al., 2022; Tuekprakhon et al., 2022). Indeed, several studies show that the Delta variant and the Omicron sub-variants are associated with significant decreases in neutralization by antibodies (Cao, Yisimayi, et al., 2022; Dejnirattisai et al., 2022; de Souza, de Freitas Amorim, Guardia, Dos Santos, Ulrich, et al., 2022; Gobeil et al., 2022; Stalls et al., 2022; Tuekprakhon et al., 2022). Interestingly, previous studies have shown that the reinfection risk with Omicron sub-variants was 6-fold higher than with other SARS-CoV-2 VOCs (Nguyen et al., 2022).

**Figure 3.**
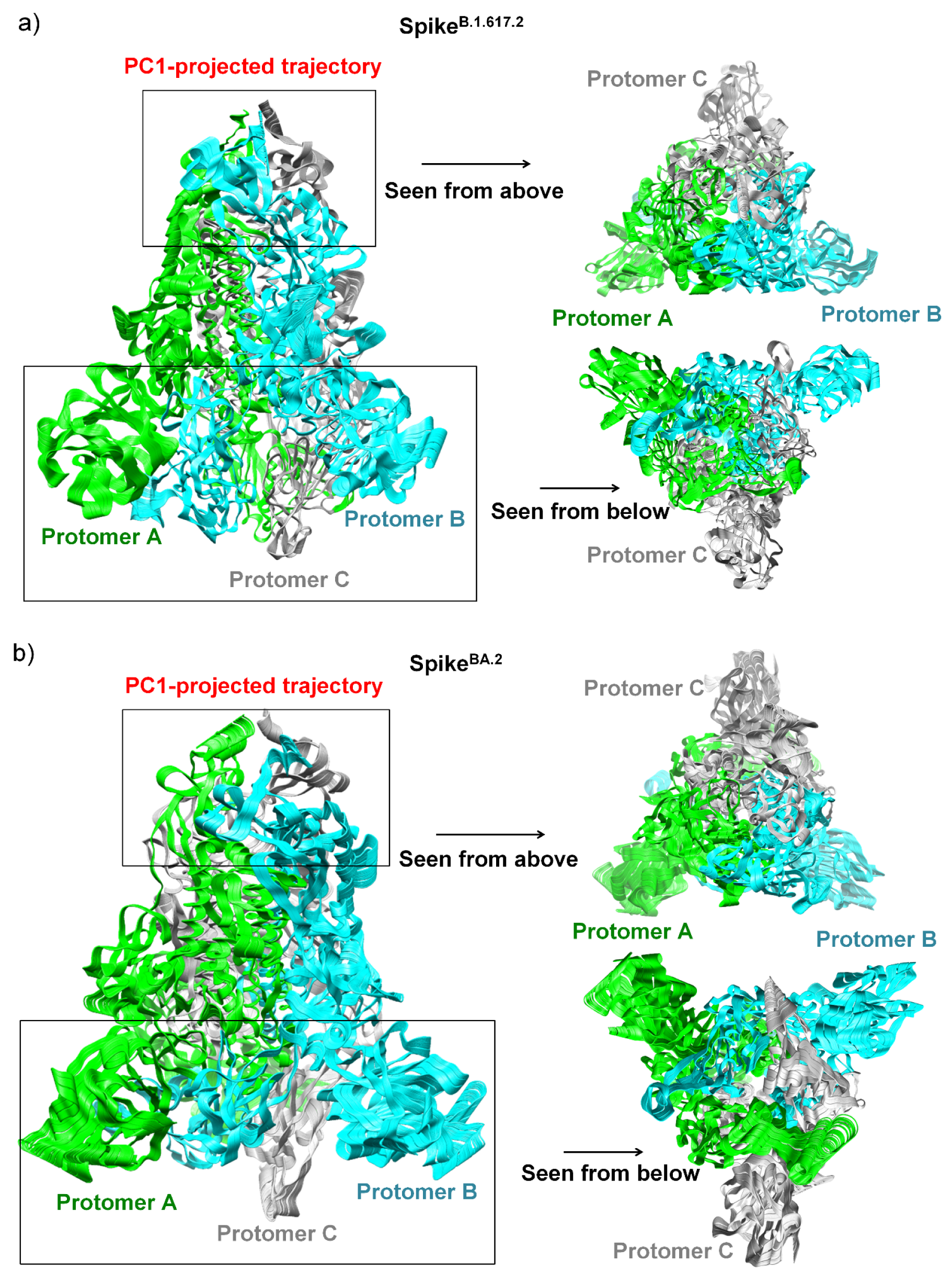
MD trajectories projected in eigenvector 1. The MD trajectory projected in eigenvector 1 (also called principal component 1, PC1) represents the main movements filtered from MD trajectory. The main movements were detected in the NTD and RBD. The panels represent structure superpositions of 100 snapshots projected in the eigenvector 1 (PC1) for **a)** Spike^B.1.617.2^ and **b)** Spike^BA.2^. Both MD projected in PC1 (skipping 300 snapshots) are available in **supplementary files S1 and S2**, which may be seen in the VMD software. The backbone conformational changes was quantified by calculating entropy using Schlitter method (ΔS^Schlitter^) at absolute temperature of 310 K. Spike^B.1.617.2^, ΔS^Schlitter^ = 74.8 kJ.mol^-1^.K^-1^; Spike^BA.2^ = 74.6 kJ.mol^-1^.K^-1^. The small rectangle is located at the S1 subunit while the big rectangle is located at the S2 subunit.

**Figure 4.**
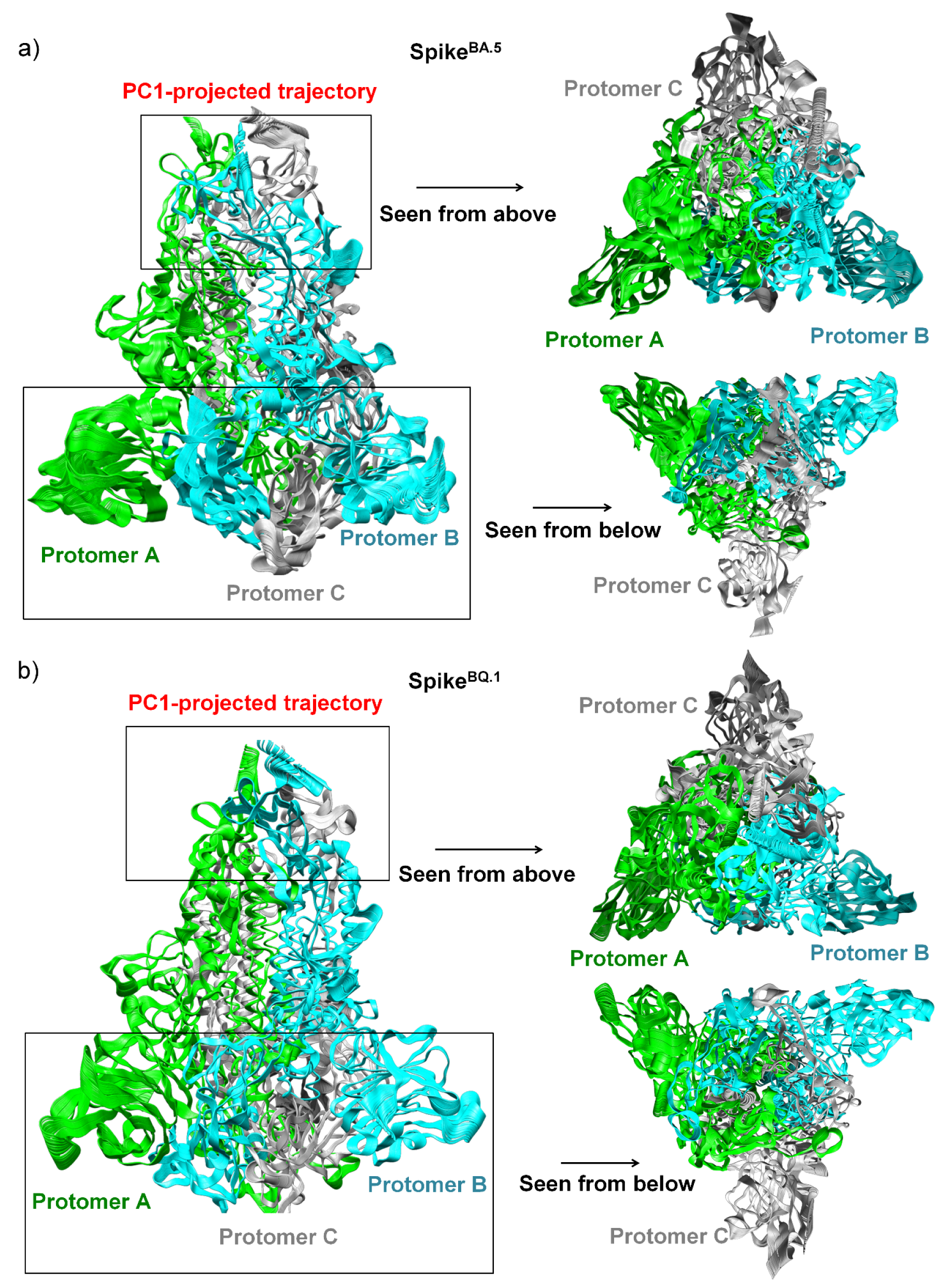
MD trajectories projected in eigenvector 1. The MD trajectory projected in eigenvector 1 (also called principal component 1, PC1) represents the main movements filtered from MD trajectory. The main movements were detected in the NTD and RBD. The panels represent structure superpositions of 100 snapshots projected in the eigenvector 1 (PC1) for **a)** Spike^BA.5^ and **b)** Spike^BQ.1^. Both MD projected in PC1 (skipping 300 snapshots) are available in **supplementary files S3 and S4**, which may be seen in the VMD software. The backbone conformational changes was quantified by calculating entropy using Schlitter method (ΔS^Schlitter^) at absolute temperature of 310 K. Spike^BA.5^, ΔS^Schlitter^ = 75.3 kJ.mol^-1^.K^-1^; Spike^BQ.1^ = 74.3 kJ.mol^-1^.K^-1^. The small rectangle is located at the S1 subunit while the big rectangle is located at the S2 subunit.

The mechanism of action of antibodies that recognize RBD is based on blocking the Spike/hACE2 interaction, preventing viral infection (Dejnirattisai et al., 2021; Yuan, Liu, et al., 2020). However, although some antibodies interact with RBD, they do not prevent interaction with hACE2 but destabilize the homotrimer (Huo et al., 2020; Yuan, Wu, et al., 2020). Taken together, our results support the hypothesis that high structural changes in NTD and RBD may significantly contribute to decreasing the action of neutralizing antibodies in immunized individuals, and may increase the transmission rate of virus.

### Mutations in K417, L452, N444, and N460 make them less exposed to interact with neutralizing antibodies

SARS-CoV-2 VOCs have several mutations that confer resistance to vaccine-induced antibodies and antibodies from convalescent patients (de Souza, de Freitas Amorim, Guardia, Dos Santos, Ulrich, et al., 2022; Tada et al., 2022; P. Wang et al., 2021). According to several studies, mutations at the sites K417, D420, and L455, all located in the RBD, may promote resistance to neutralizing antibodies (Cao, Wang, et al., 2022; Ferreira et al., 2021). In addition, in BQ.1 and BQ.1.1, the substitutions K444T and N460K favor resistance to neutralizing antibodies (Cao, Wang, et al., 2022). Since these mutations involve substitutions of amino acid residues that may make novel hydrogen bonding interactions, we investigated how these substitutions could affect the recognition of neutralizing antibodies due to increased interactions with local amino acid residues. To verify this hypothesis, we calculated the hydrogen bonding occupations of Spike^B.1.617.2^, Spike^BA.2^, Spike^BA.5^, and Spike^BQ.1^. Hereafter, occupancies are expressed as the proportion of simulation frames where the residues are close enough and in a suitable geometry for the formation of hydrogen bonding interactions.

In order to investigate the effects of substitutions K417N, L452R, K444T, and N460K in the hydrogen bonding networks, we calculated the hydrogen bonding occupancies (**Tables S1-S4**). In Spike^B.1.617.2^, residue K417^B.1.617.2^ side chain (SC) interacts with _SC_N422^B.1.617.2^ (occupancy = 35.96%), _SC_E406^B.1.617.2^ (76.32%), _SC_Y421^B.1.617.2^ (27.40%), and main chain (MC) of _MC_F374^B.1.617.2^ (30.24%). In Spike^BA.2^, the mutation K417N changed the interactions of the corresponding residue (_SC_N414^BA.2^) by adding hydrogen bonding interactions with _SC_N367^BA.2^ (33.41%) and _MC_R451^BA.2^ (62.75%). In Spike^BA.5^, we observed that _SC_N412^BA.5^ interacts with _SC_Q404^BA.5^ (17.37%), _MC_R449^BA.5^ (66.12%), and _SC_Y416^BA.5^ (10.72%). For Spike^BQ.1^, the equivalent residue, _SC_N411^BQ.1^ presents increased hydrogen bonding interaction networks with residues _SC_E409^BQ.1^ (28.68%), _MC_A375^BQ.1^ (15.03%), _MC_Y372^BQ.1^ (23.51%), _MC_R457^BQ.1^ (49.13%), _MC_L458^BQ.1^ (10.31%), and _SC_Y424^BQ.1^ (12.36%). Overall, these results demonstrate a global increase of hydrogen bonding interactions and a more complex network of interactions between local residues. The K417N mutation has been demonstrated to be critical to evade the immune system (Hirabara et al., 2021). Therefore, the K417N substitution expands the network of interactions to neighboring residues, possibly making this asparagine residue less exposed to interact with neutralizing antibodies.

We also investigated how the L452R substitution influences neutralizing antibody resistance. In Spike^B.1.617.2^, we observed that _SC_R452^B.1.617.2^ makes hydrogen bonding interactions with residues _SC_Y351^B.1.617.2^ (18.41%), _MC_N450^B.1.617.2^ (23.53%), _SC_E484^B.1.617.2^ (87.43%), and _SC_S494^SC^ (9.43%). This network of interactions may be directly related to the loss of binding affinity with neutralizing antibodies, where _SC_R452^B.1.617.2^ may preferentially interact with these amino acids, weakening interactions with neutralizing antibodies. On the other hand, in Spike^BA.2^, _SC_L452^BA.2^ residue does not mutate and thus preserved hydrophobic interactions with residues _SC_Y348^BA.2^, _SC_Y448^BA.2^, _SC_L449^BA.2^, _SC_Y450^BA.2^, _SC_C477^BA.2^, _SC_G479^BA.2^, _SC_V480^BA.2^, _SC_A481^BA.2^, _SC_Y486^BA.2^, _SC_F487^BA.2^, _SC_L489^BA.2^. In Spike^BA.5^, _SC_R447^BA.5^ interacts with residues _SC_Y346^BA.5^ (35.37%), _MC_N445^BA.5^ (29.54%), _SC_S489^BA.5^ (18.39%), and _MC_S489^BA.5^ (14.23%). In Spike^BQ.1^, a wide network of hydrogen bonding interactions forms between _SC_R445^BQ.1^ and residues _SC_D436^BQ.1^ (86.92%), _MC_S437^BQ.1^ (18.56%), _SC_F341^BQ.1^ (35.88%), _SC_N444^BQ.1^ (43.39%), _SC_Y489^BQ.1^ (20.50%), and _MC_Y489^BQ.1^ (16.10%). L452R substitution has been reported to escape HLA-A24-restricted cellular immunity (Motozono et al., 2021). Therefore, the Spike^BQ.1^ contains an increase in hydrogen bonding interactions due to the L452R substitution. Thus, our results show that the mutation L452R expands the hydrogen bonding interactions with neighboring residues, a change that may likely reduce the availability of this residue for interactions with neutralizing antibodies against SARS-CoV-2.

BQ.1 and BQ.1.1 also have two other substitutions, K444T and N460K, that are known to favor resistance to neutralizing antibodies (Cao, Wang, et al., 2022). We measured the hydrogen bonding interaction occupancies for residues _SC_K439^BA.5^ and _SC_T438^BQ.1^ with neighboring residues. Residue _SC_K439^BA.5^ makes hydrogen bonding interactions with _MC_N443^BA.5^ (11.29%), _MC_G442^BA.5^ (22.57%), and _SC_N443^BA.5^ (10.56%). However, _SC_T438^BQ.1^ interacts with _MC_S432^BQ.1^ (14.12%), _MC_D436^BQ.1^ (12.49%), _SC_N442^BQ.1^ (43.39%), and _SC_R503^BQ.1^ (53.53%). Thus, the N444T mutation (_SC_T438^BQ.1^) probably also leads to an escape of the immune system recognition, as suggested from a significant increase in hydrogen bonding occupancy for this threonine residue. Most notably, a large change in occupancy for the hydrogen bonding interaction between _SC_T438^BQ.1^ and _SC_N442^BQ.1^ and the emergency of interactions with _SC_R503^BQ.1^ highlights the differences between BA.5 and BQ.1. In addition, in Spike^B.1.617.2^, hydrogen bonding interactions are observed between _SC_N460^B.1.617.2^ and the residues _SC_D420^B.1.617.2^ (80.47%), _SC_K424^B.1.617.2^ (62.39%), and _SC_Y421^B.1.617.2^ (39.35%). For Spike^BA.2^, there are interactions between _SC_N457^BA.2^ and _SC_D417^BA.2^ (74.96%), and _SC_K421^BA.2^ (63.93%). Spike^BA.5^ presents hydrogen bonding interactions between _SC_N455^BA.5^ and residues _SC_K419^BA.5^ (38.71%), _SC_D415^BA.5^ (57.35%) and _SC_Y416^BA.5^ (10.11%). For Spike^BQ.1^, interactions are observed between _SC_K454^BQ.1^ and _SC_D979^BQ.1^ (35.07%), _SC_D414^BQ.1^ (94.89%) and _SC_Y415^BQ.1^ (38.01%).

Our results suggest SARS-CoV-2 Omicron sub-variants are evolving through the acquisition of evolutionary advantageous mutations that enhance interactions of T438^BQ.1^ and residues N442^BQ.1^ and R503^BQ.1^, thus decreasing the availability of these amino acids to neutralizing antibodies. Another fundamental contribution is the N460K substitution observed in Spike^BQ.1^, which mediates a strong saline interaction between _SC_K454^BQ.1^ and _SC_D414^BQ.1^ and thus may also contribute to a low availability of the _SC_K454^BQ.1^ for interactions with neutralizing antibodies. Indeed, Qu and co-authors showed that K444T and N460K mutations in Spike^BQ.1^ and Spike^BQ.1.1^ promote resistance to neutralizing antibodies (Qu et al., 2022). In addition, previous studies have reported that people previously infected with Omicron can be reinfected with different Omicron sub-variants in a shorter time than SARS-CoV-2 VOCs. In this case, the reinfection can happen after two months (Nguyen et al., 2022). Thus, our results are important because they show how the Spike protein evolves into variants that hide the recognition region of neutralizing antibodies, increasing immune system evasion and consequently transmission of SARS-CoV-2.

### Cooperative behavior of D614G substitution in Spike^B.1.617.2^, Spike^BA.2^, Spike^BA.5^ and Spike^BQ.1^

According to cryo-EM studies, the D614G substitution does not cause high conformational changes in the Spike structure, except for the loss of the hydrogen bonding interaction between D614^WT^ and K854^WT^, favoring the exposure of RBD to interact with hACE2 (J. Zhang et al., 2021). Our previous study has shown that D614G is predominant in the Spikes of SARS-CoV-2 VOCs (de Souza, de Freitas Amorim, Guardia, Dos Santos, Ulrich, et al., 2022). This mutation also is essential for the virus because **(1)** it causes the disruption of the interprotomer contact, increasing up-RBD state (Mansbach et al., 2021b); **(2)** modulates structural conformations that promote membrane fusion (J. Zhang et al., 2021); **(3)** increases virion spike density and infectivity (Yurkovetskiy et al., 2020; L. Zhang et al., 2020); and **(4)** increases the cleavage rate by furin (Gobeil et al., 2021). Our previous molecular dynamics study of the Spike^BA.1^ has shown that D614G mutation contributes to exposure RBD due to loss of hydrogen bonding interactions between _SC_D614^BA.1^ the residue pairs _SC_K854, _SC_Y837^BA.1^, _SC_K835^BA.1^, _SC_T859^BA.1^ (de Souza, Amorim, et al., 2022). In order to understand the impact of the D614G mutation in the Spike^B.1.617.2^, Spike^BA.2^, Spike^BA.5^, and Spike^BQ.1^, we calculated the dynamical cross-correlation matrix (DCCM) between all residue pairs *i* and *j* using molecular dynamics trajectories. DCCM is defined as the matrix *DCC_ij_* = ⟨Δ*R_i_*. Δ*R_j_*, where Δ*R_i_* = *R_i_* − ⟨*R_j_*⟩ is the difference between the Euclidean vector R_i_, for position of the C_α_ of residue *i*, and the average position of this residue, ⟨R_i_⟩. Next, we calculated the value for the Pearson correlation matrix after normalization as:

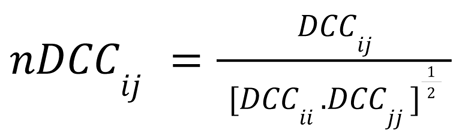

where *nDCC_ij_* can range between −1 and 1. The *nDCC_ij_* = 0 means no correlations, *nDCC_ij_* = 1 indicates complete correlation and *nDCC_ij_* =− 1 shows complete anticorrelation. We used DCCM for investigating cooperativity between protomores within the trimeric structure of the Spike. The complete correlation matrices are available in the **supplementary files S5-S8**. Due to the high number of uncorrelated residue pairs, we applied a cut-off value of 0.6 to the nDCC_ij_ values (**supplementary files S9-S12**). We then focused only on the correlations between residues equivalent to D614^WT^ (G614^B.1.617.2^, G611^BA.2^, G609^BA.5^ and G608^BQ.1^) and all other amino acid residues (**Figures 5-8**). For all Spike sub-variants, intra-protomer correlations between residues equivalent to D614^WT^ and P681^WT^ and their adjacent residues correspond to local hydrogen bonding interactions. In Spike^B.1.617.2^, we observed, for the protomers B and C, an intra-protomer correlation between G614 and the residues 316-319 (region between NTD and RBD) (**Figure 5**). In contrast, for the Spike^BA.2^, both intra- and inter-protomer correlations were observed. Spike^BA.2^ has correlations between G611_C_ and residues G32-V33_A_, N162_A_, and D284-V286_A_, all of which are located in the NTD, the subscripts correspond to the A, B or C chains of trimeric Spike structure (**Figure 6**). However, Spike^BA.2^ also shows intra-protomer correlations for the G611 residue of all chains and residues located between the NTD and RBD (S313-V317) and anticorrelations for residues A369 and P488 (**Figure 6**). In Spike^BA.5^, there is an intra-protomer correlation of the G609_A_ with residues between NTD and RBD (F313-S320_A_) and between G609_C_ and residues I307-Q309_C_ and S313_C_ (**Figure 7**). Spike^BQ.1^ also showed only two intra-protomer correlations with the region between NTD and RBD, involving residue G608_A_ and residues N311-V314_A_ and residue G608_B_ and F312_B_ (**Figure 8**). In our previous study, Spike^WT^ showed few cooperative relationships, including a correlation of D614_B_ and the residues S71_A_ (in NTD) and I312_B_, which is located between NTD and RBD (de Souza, Amorim, et al., 2022). On the other hand, the D614G mutation in Spike^BA.1^ gave origin to a direct communication between G611^BA.1^ and residues belonging to NTD and between NTD and RBD (de Souza, Amorim, et al., 2022). Overall, the D614G substitution seems to play a key role in the emergency of communication between D614G with residues in the β-sheet located between the NTD and the RBD (residues 306-334) (Gobeil et al., 2021). Taken together, our results are in agreement with previous studies (Gobeil et al., 2021), where small structural changes in this region could be critical to generate significant cooperative effects that will interfere with the availability of RBD for the interaction with hACE2 or affect the efficiency of neutralizing antibodies.

**Figure 5.**
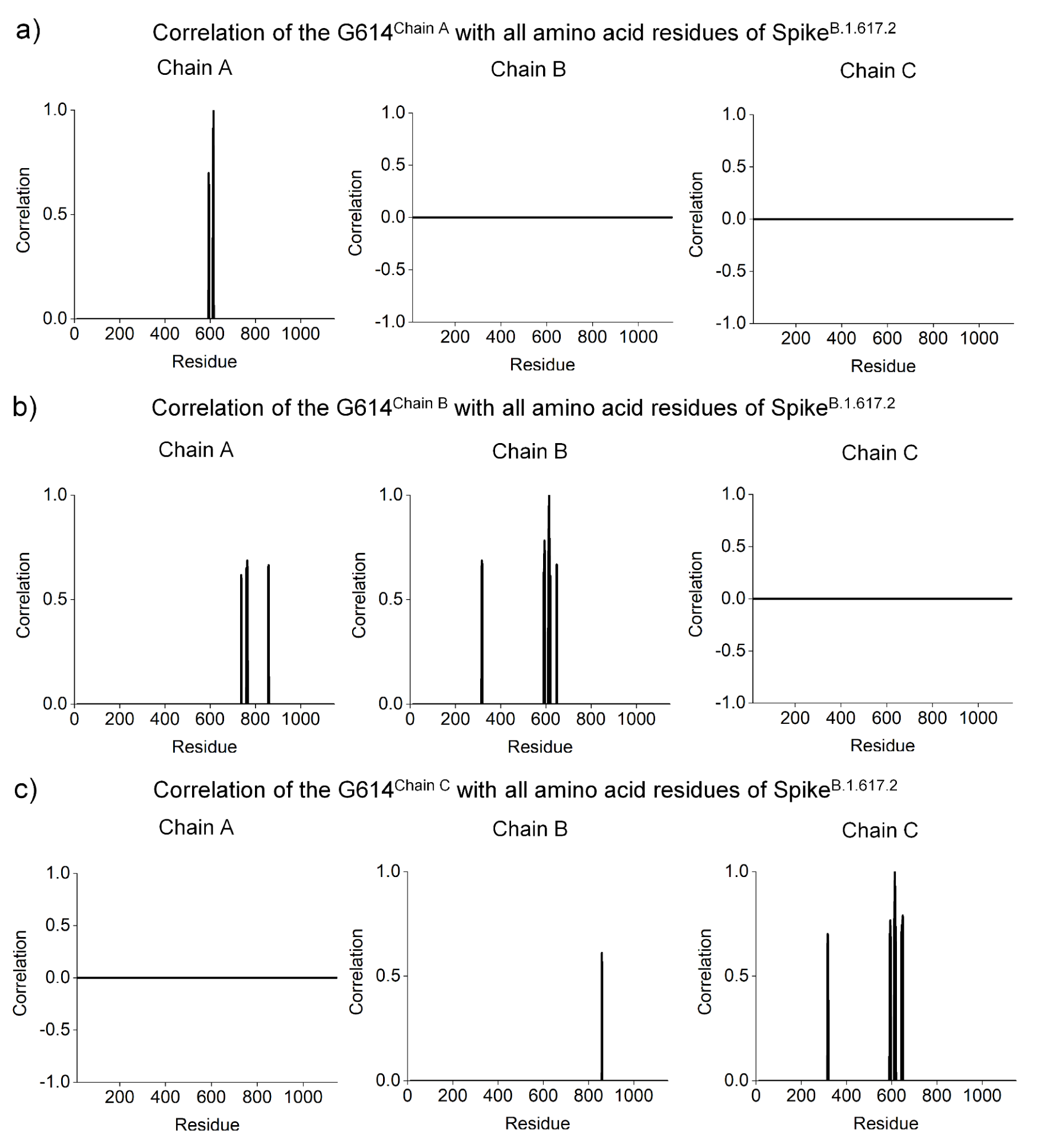
Dynamical cross-correlation matrix analysis for investigating the cooperative between protomers within the trimeric form of the Spike structure affected by the D614G substitution in the Spike^Delta^. Correlations up to 0.6 and below -0.6 are shown. **a)** G614^A^ with all amino acid residues of the protomers A, B and C. The results are [G614^A^ x residues^A^ (593-595, 612-615)]; **b)** G614^B^ with all amino acid residues of the protomers A, B and C [G614^B^ x residues^A^ (737-738, 760-761, 763-765, 857-859), G614^B^ x residues^B^ (316-319, 591-595, 611-617, 620, 648-649)].; **c)** G614^C^ with amino acid residues of the protomers A, B and C [G614 ^C^ x residues^B^ (859), G614^C^ x residues^C^ (316-319, 591-596, 611-618, 643-650)].

**Figure 6.**
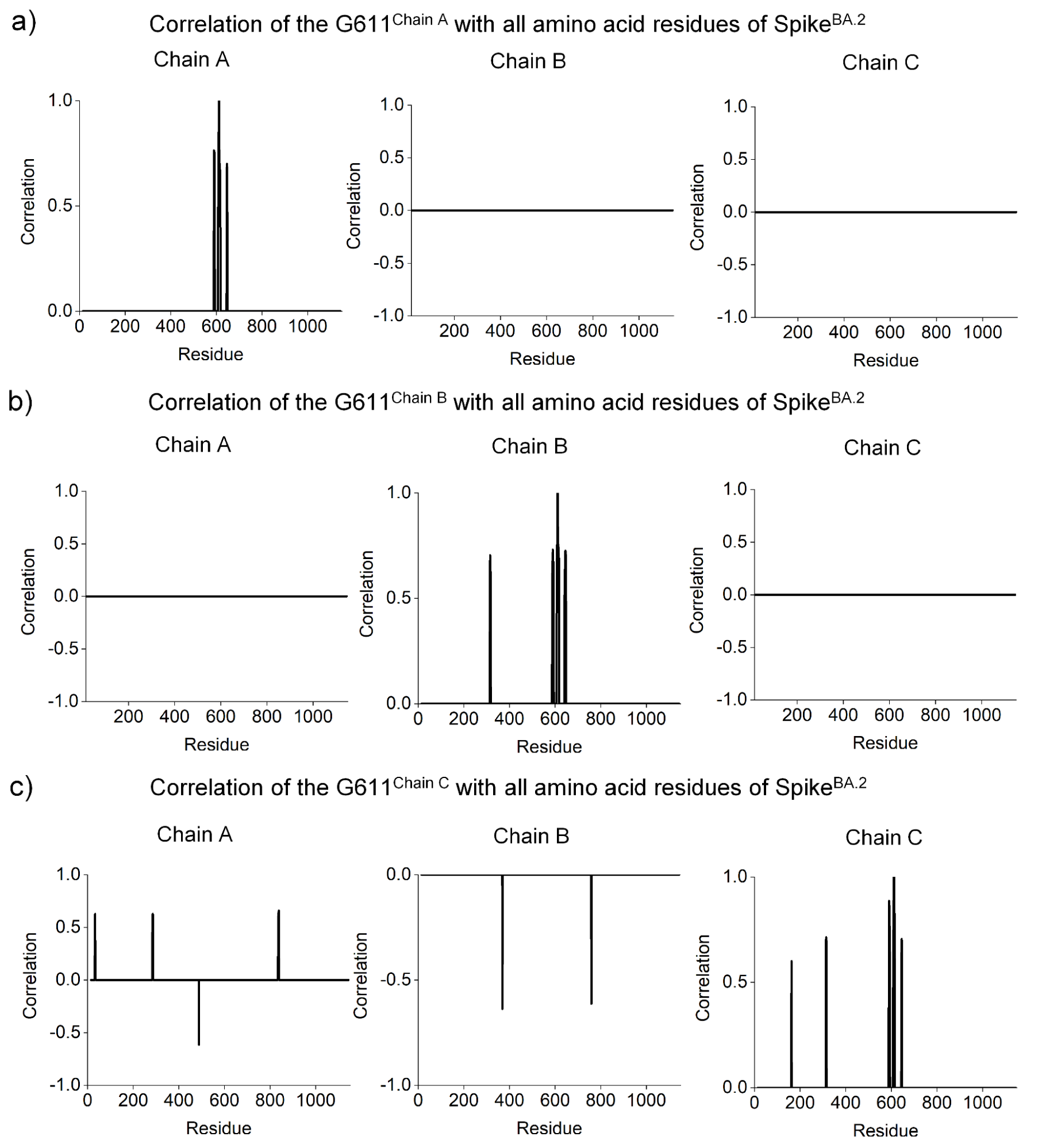
Correlation analysis for investigating the cooperative effect of the D614G substitution in the Spike^BA.2^. Correlations up to 0.6 and below -0.6 are shown. **a)** G611^A^ with amino acid residues of the protomers A, B and C [G611^A^ x residues^A^ (589-592, 608-617, 645-647)]; **b)** G611^B^ with amino acid residues of the protomers A, B and C [G611^B^ x residues^B^ (314-317, 587-593, 607-617, 641-647)]; **c)** G611^ChainC^ with amino acid residues of the protomers A, B and C [G611^C^ x residues^A^ (32-33, 284-286, 488, 835-838), G611^C^ x residues^B^ (369, 759), G611^C^ x residues^C^ (162, 313-315, 589-593, 608-614, 644-646)].

**Figure 7.**
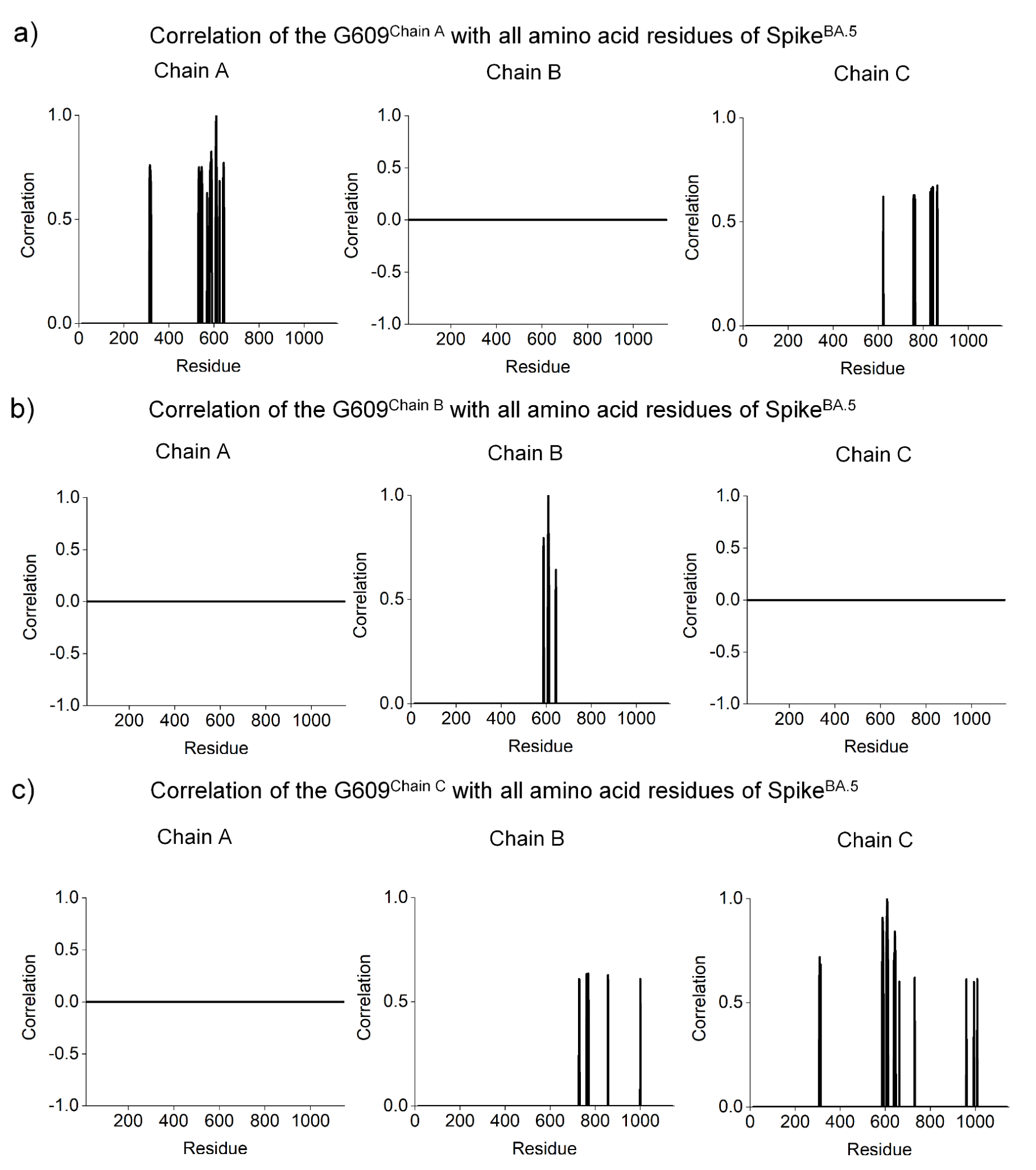
Correlation analysis for investigating the cooperative effect of the D614G substitution in the Spike^BA.5^. Correlations up to 0.6 and below -0.6 are shown. **a)** G609^A^ with amino acid residues of the protomers A, B and C [G609^A^ x residues^A^ (313-320, 531-536, 543-547, 569, 581-590, 607-612, 624, 640-644), G609^A^ x residues^C^ (622, 756-757, 760-761, 763, 831-835, 842-843, 861-862)]; **b)** G609^B^ with amino acid residues of the protomers A, B and C [G609^B^ x residues^B^ (587-589, 607-612, 642-643)]; **c)** G609^C^ with amino acid residues of the protomers A, B and C [G609^C^ x residues^B^ (728, 730, 761-762, 765, 769-770, 856-857, 1000), G609^C^ x residues^C^ (307-309, 313, 587-592, 605-613, 639-646, 663, 731, 960-961, 994)].

**Figure 8.**
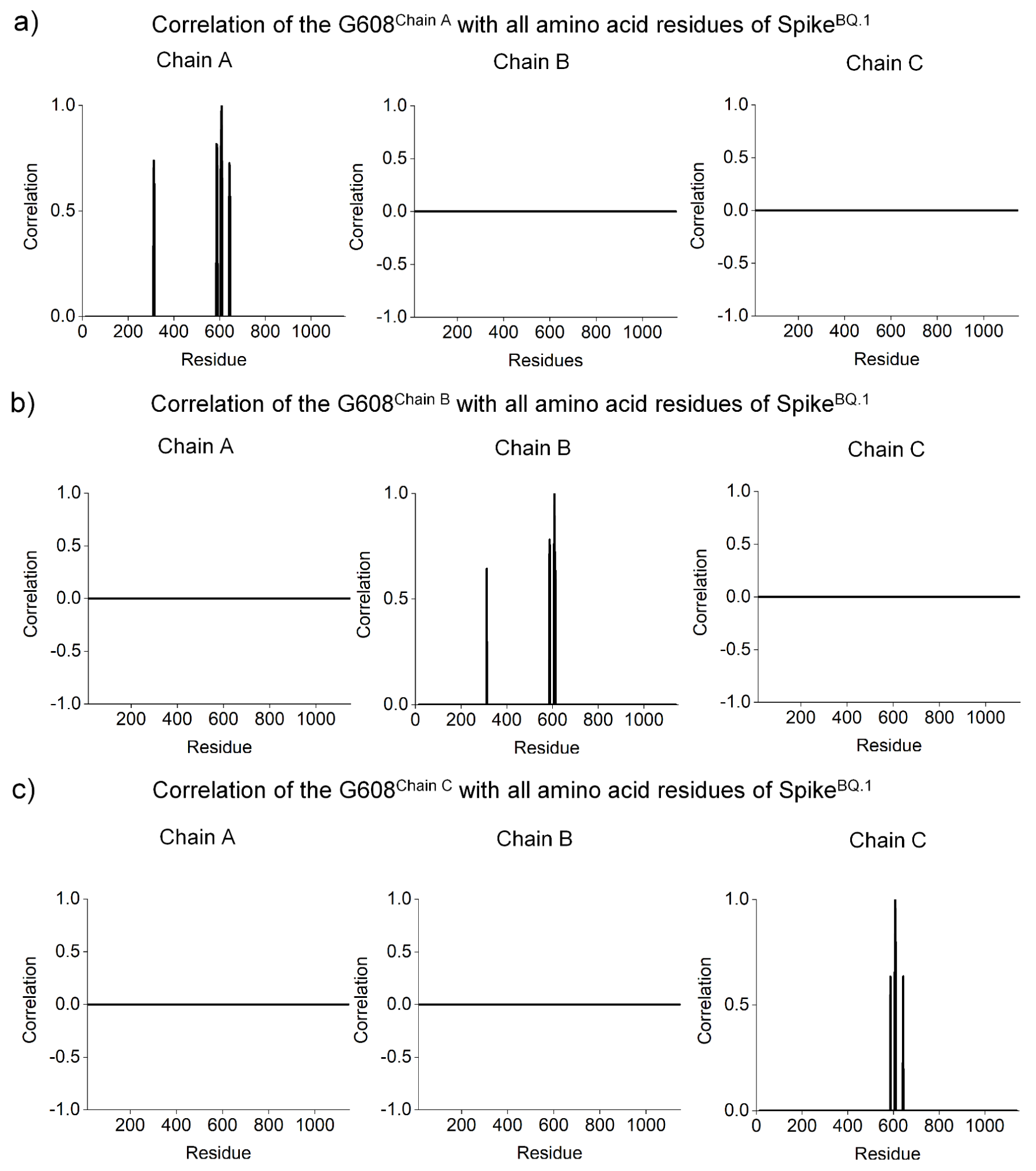
Correlation analysis for investigating the cooperative effect of the D614G substitution in the Spike^BQ.1^. Correlations up to 0.6 and below -0.6 are shown. **a)** G608^A^ with amino acid residues of the protomers A, B and C [G608^A^ x residues^A^ (311-314, 586-590, 604-611, 642-645)]; **b)** G608^B^ with amino acid residues of the protomers A, B and C [G608^B^ x residues^B^ (312, 586-588, 606-611, 613)]; **c)** G608^C^ with amino acid residues of the protomers A, B and C [G608^C^ x residues^C^ (587-588, 606-610, 642-643)].

### Increased cooperativity supports a role of P681R mutation in high lethality the SARS-CoV-2 Delta Variant

In **Figure S2**, we show that all Omicron sub-variants carry a P681H substitution while the Delta variant has a P681R substitution. In order to understand the effect of these substitutions in the evolution of the Delta and the Omicron sub-variants, we investigated the impact of these substitutions in the network of correlated residues. The detailed results of the DCCM analysis for the substitutions P681R in Spike^B.1.617.2^ and P681H in Spike^BA.2^, Spike^BA.5^, and Spike^BQ.1^ are shown in **Figures 9-12**. In Spike^B.1.617.2^, the P681R substitution causes a significant gain of correlations between distant residues, thus implying higher intra- and inter-protomer communication between S1 and S2 subunits than in the wild type (**Figure 9**). In contrast, we clearly noted that the P681H substitutions of the Omicron sub-variants tend to decrease the inter-protomer correlations between the S1 and S2 subunits. A previous study showed that Spike^B.1.1.539^ is significantly less fusogenic than the Spike^B.1.617.2^ (Suzuki et al., 2022). In addition, Suzuki and co-authors also showed that hamsters infected with SARS-CoV-2^Omicron^ exhibited fewer respiratory disturbances than those infected with SARS-CoV-2^B.1.617.2^, suggesting that Omicron is less virulent than Delta (Suzuki et al., 2022). Our results suggest that the P681R mutation favors intra- and inter-protomer communication between S1 and S2 subunits. Therefore, given that Omicron is less fusogenic than Delta, we hypothesize that the abundant long-distance communication may be a key factor to the higher fusogenicity of the Delta variant when compared to Omicron sub-variants.

**Figure 9.**
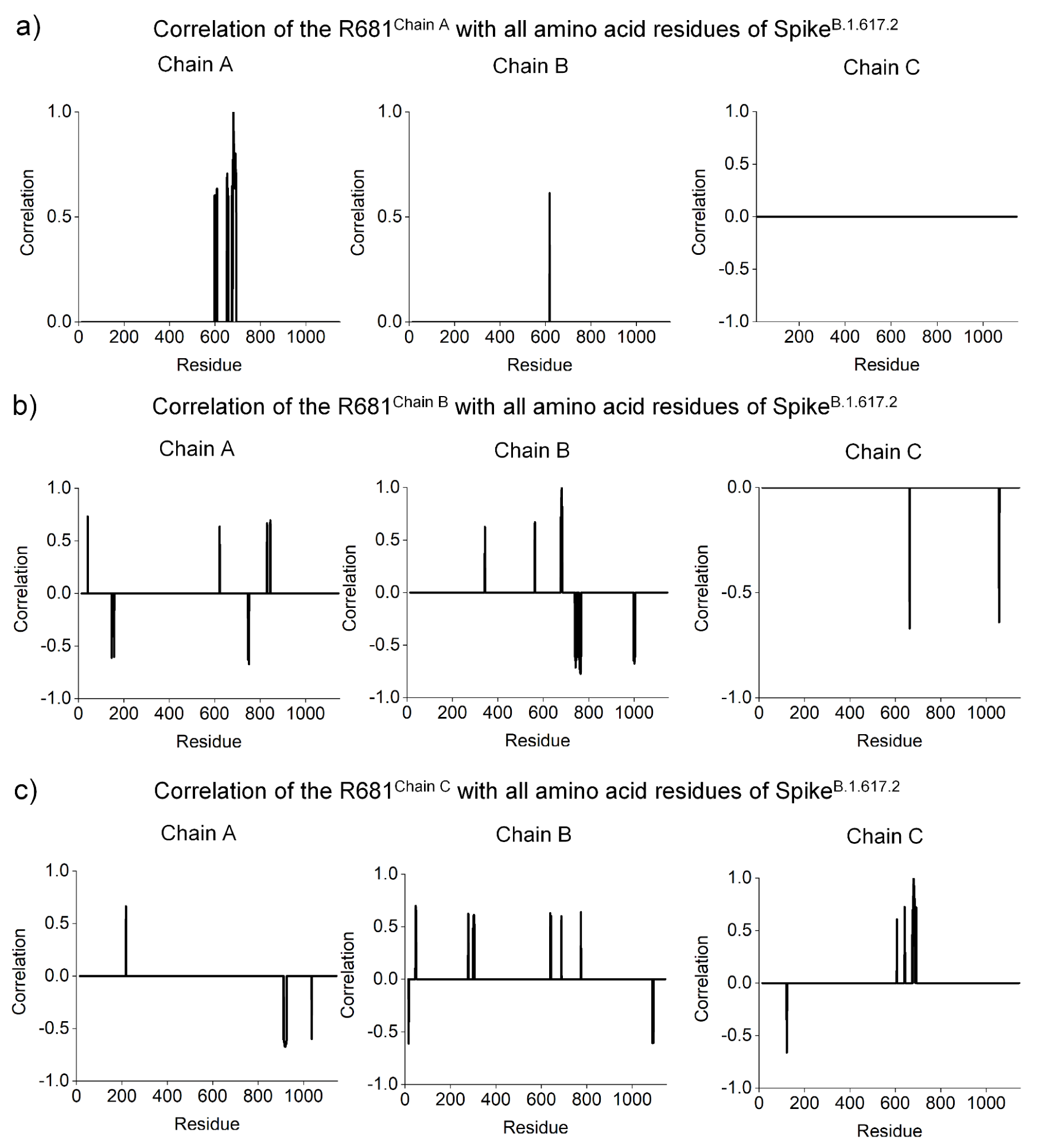
Correlation analysis for investigating the cooperative effect of the P681R substitution in the Spike^B.1.617.2^. Correlations up to 0.6 and below -0.6 are shown. **a)** R681^A^ with all amino acid residues of the protomers A, B and C [R681^A^ x residues^A^ (598-600, 607-609, 652-654, 656, 658, 674-676, 678-693), R681^A^ x residues^B^ (618)]; **b)** R681^B^ with all amino acid residues of the protomers A, B and C [R681^B^ x residues^A^ (41, 146, 157, 622, 747, 750-751, 830-831, 844-846), R681^B^ x residues^B^ (343, 562-563, 677-683, 739, 741-745, 753, 758-766, 997, 1000-1004), R681^B^ x residues^C^ (663, 1057-1058)]; **c)** R681^C^ with amino acid residues of the protomers A, B and C [R681^C^ x residues^A^ (218, 912, 914-925, 1036), R681^C^ x residues^B^ (16, 46-49, 278-279, 300, 303, 305, 639, 642, 688, 774, 1089, 1093), R681^C^ x residues^C^ (122-123, 607, 641-642, 675, 678-687, 690, 692)].

In contrast, it is already known that the Omicron variant spreads faster than Delta (Nishiura et al., 2021) and is more resistant to neutralizing antibodies than other SARS-CoV-2 VOCs, including Delta (Cameroni et al., 2022; Cao, Wang, et al., 2022; Cele et al., 2022; Liu et al., 2022; Meng et al., 2022). Conversely, all Omicron sub-variants are highly transmissible and carry the P681H substitution (in Spike^B.1.1.539^) (Suzuki et al., 2022). Although still unclear, the low lethality of the Omicron sub-variants may be explained in part due to the decreased virulence of Omicron, which might be in part a consequence of the wide coverage of the global vaccination rate in humans (C. Wang et al., 2022). In our computational analysis, a decrease in the cooperative behavior is strongly associated with the P681H substitution and probably affects Spike function via impairment of protomer-protomer interactions (**Figures 10-12**). We propose that the low lethality of Omicron sub-variants and the associated decrease of virus fusogenicity could be explained by the loss of cooperativity caused by mutations such as the P681H substitution. In contrast, the P681R substitution present in the Delta variant is correlated with an elevated number of intra- and inter-domain correlations and, critically, this mutation contributes to enhanced communication among subdomains responsible for initiation of the hACE2 interactions and those associated with membrane fusion in a mechanism not well described. P681R, therefore, probably contributes greatly to enhancing fusogenicity, thus leading to higher infectivity and lethality.

**Figure 10.**
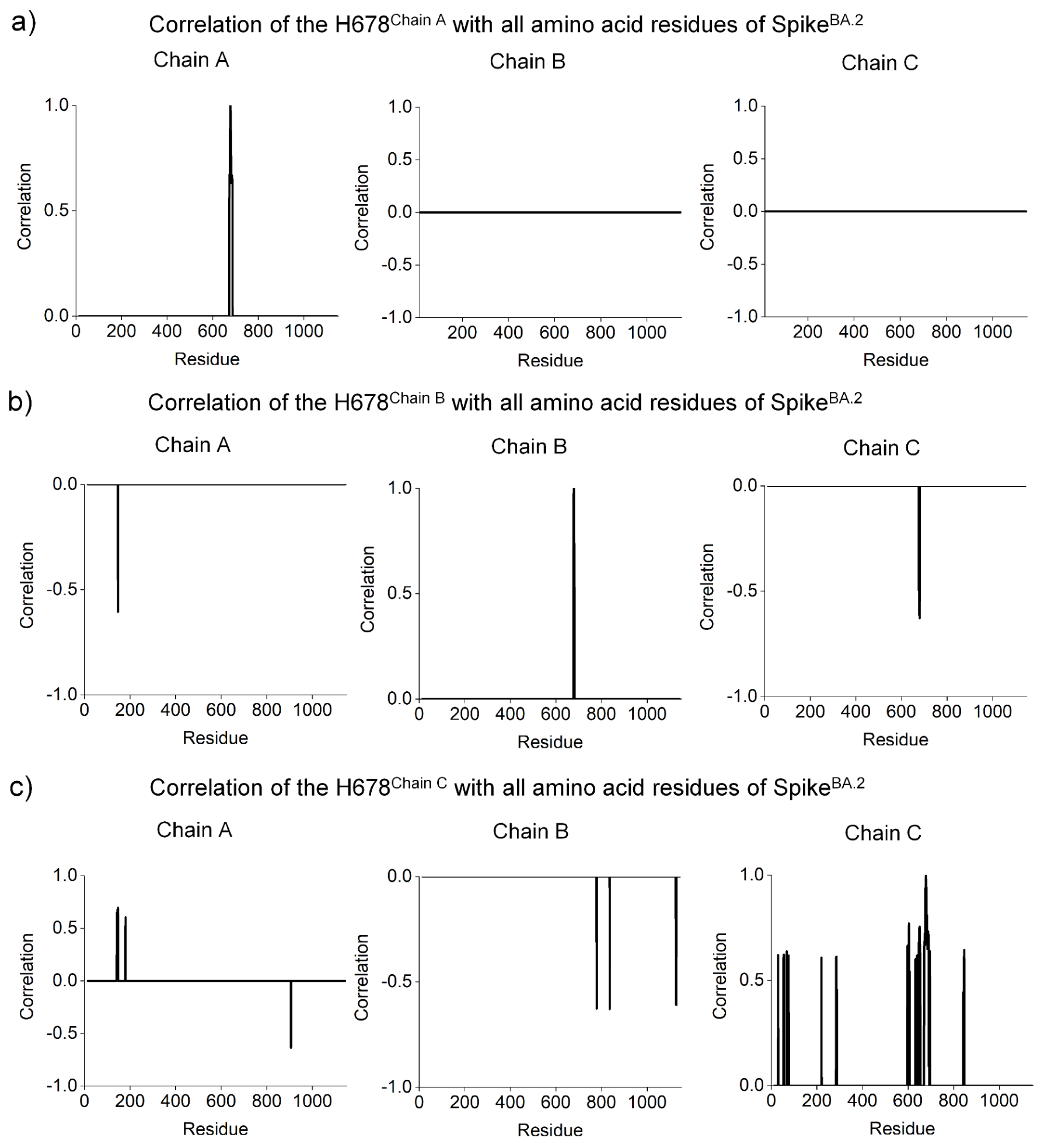
Correlation analysis for investigating the cooperative effect of the P681H substitution in the Spike^BA.2^. Correlations up to 0.6 and below -0.6 between are shown. **a)** H678^ChainA^ with all amino acid residues of the protomers A, B and C [H678^A^ x residues^A^ (673-686)]; **b)** R678^ChainB^ with all amino acid residues of the protomers A, B and C [H678^B^ x residues^A^ (147), H678^B^ x residues^B^ (677-681), H678^B^ x residues^C^ (677, 680)]; **c)** R678^ChainC^ with amino acid residues of the protomers A, B and C [H678^C^ x residues^A^ (142-145, 147-148, 180, 906-907), H678^C^ x residues^B^ (779, 836, 1127-1128), H678^C^ x residues^C^ (30, 54-55, 68, 76, 220, 285-287, 598-599, 601, 603-606, 633, 639-640, 648-654, 671-691, 696, 844-845, 847)].

**Figure 11.**
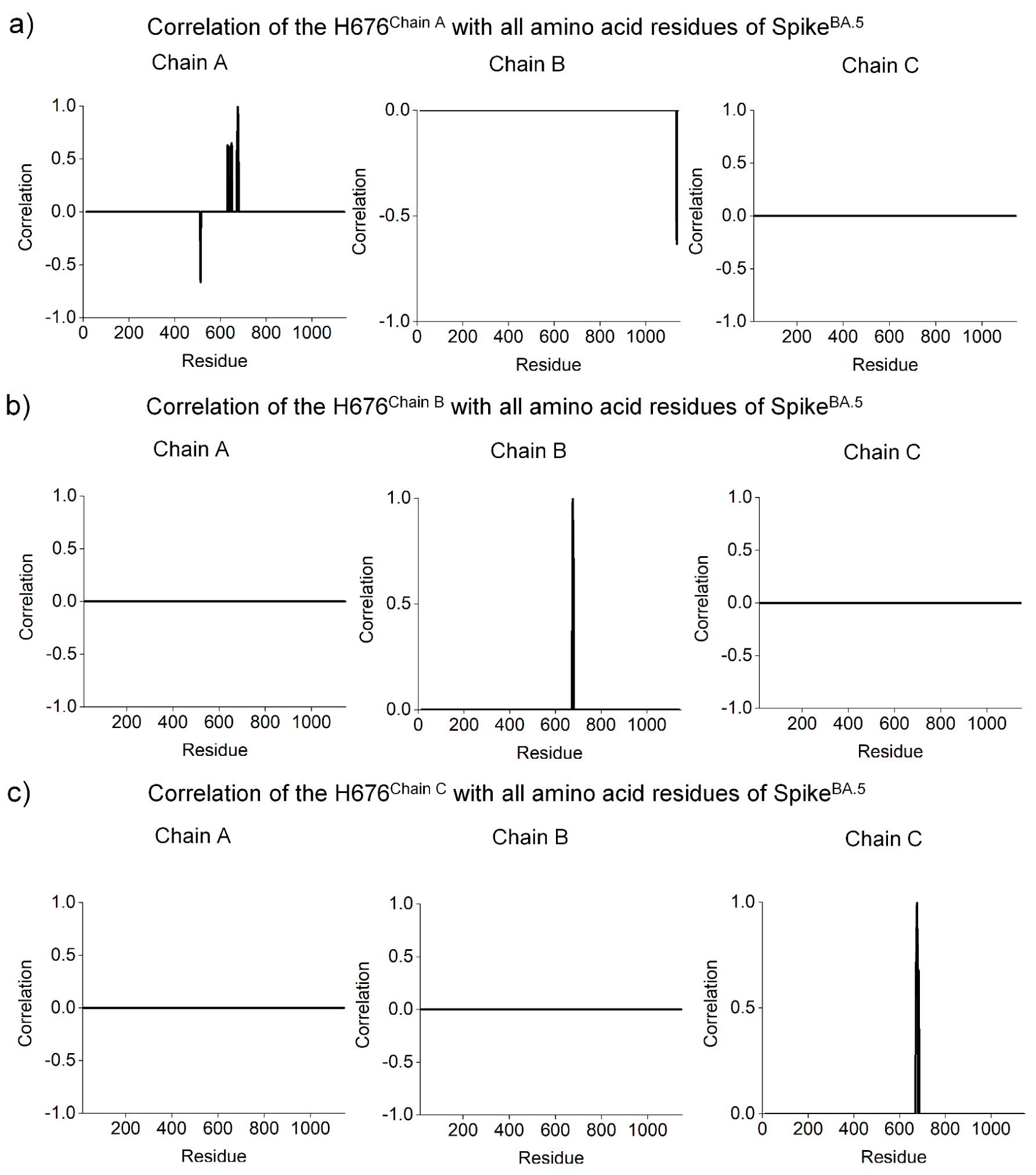
Correlation analysis for investigating the cooperative effect of the P681H substitution in the Spike^BA.5^. Correlations up to 0.6 and below -0.6 are shown. **a)** H676^A^ with all amino acid residues of the protomers A, B and C [H676^A^ x residues^A^ (512-514, 631, 633, 635, 646-650, 672-679)]; **b)** R676^B^ with all amino acid residues of the protomers A, B and C [H676^B^ x residues^A^ (1134-1136), H676^B^ x residues^B^ (674-680)]; **c)** R676^C^ with amino acid residues of the protomers A, B and C [H676^C^ x residues^C^ (671-681, 684-685)].

**Figure 12.**
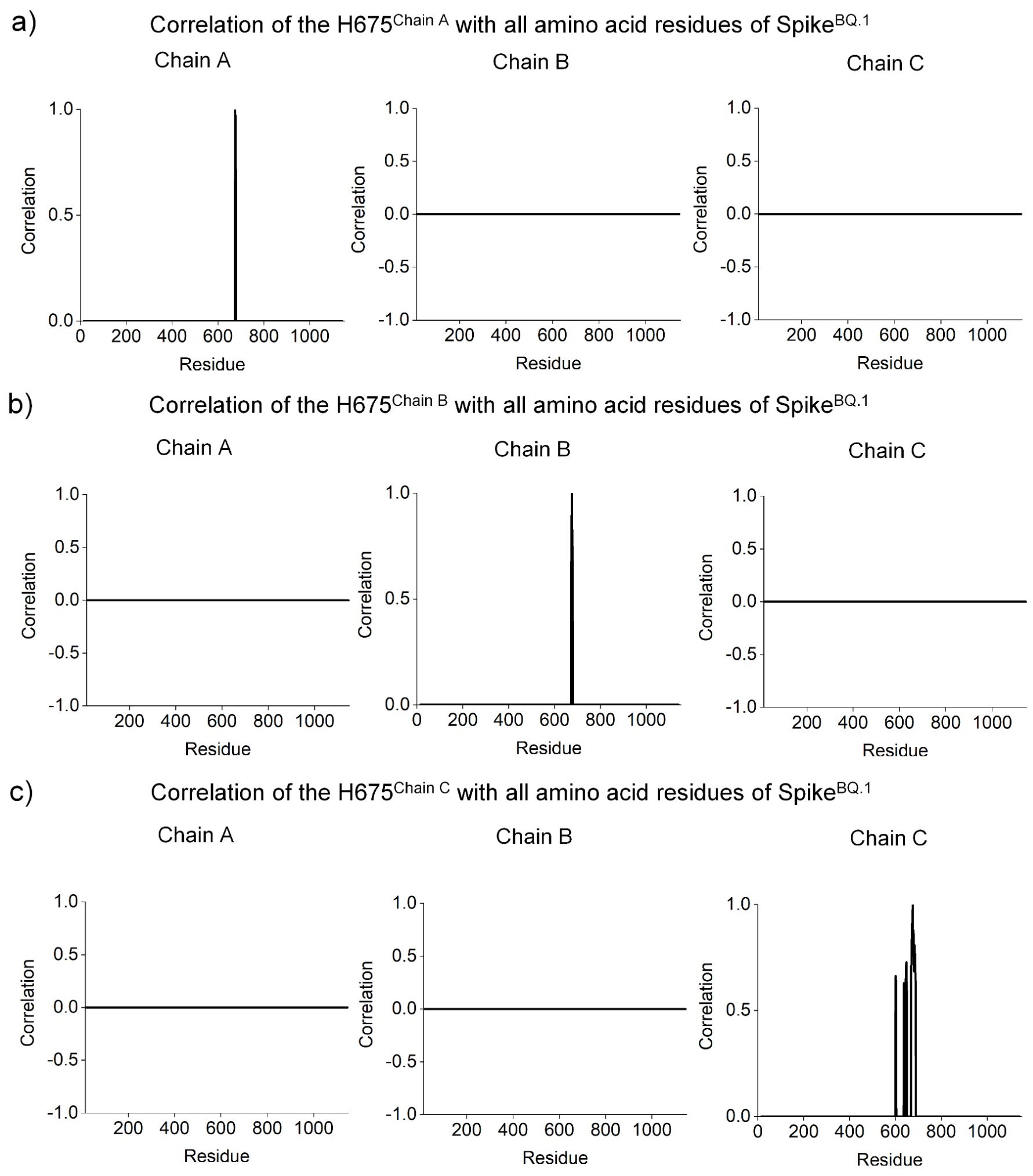
Correlation analysis for investigating the cooperative effect of the P681H substitution in the Spike^BQ.1^. Correlations up to 0.6 and below -0.6 are shown. **a)** H675^A^ with all amino acid residues of the protomers A, B and C [H675^A^ x residues^A^ (674-678)]; **b)** H675^B^ with all amino acid residues of the protomers A, B and C [H675^B^ x residues^B^ (673-679)]; **c)** H675^C^ with amino acid residues of the protomers A, B and C [H675^C^ x residues^C^ (600-603, 637, 650, 669-688)].

## Conclusion

The degree of virulence of SARS-CoV-2 is dependent on the genomic mutations carried by the virus, which impact process such recognition by hACE2, cleavage rate by proteases (furins, TMPRSS2 and cathepsins), and the fusogenic processes. In addition, transmissibility is also directly related to immune system escape and, together with pathogenicity, impacts hospitalizations, severe cases and deaths. The Omicron variant is known to be less virulent than WT or other previous VOCs (Meng et al., 2022; Peacock et al., 2022; Shuai et al., 2022; Yamamoto et al., 2022). BA.5 and BQ.1 are predominant worldwide (including B.1.617.2 and BA.2), which motivated us to investigate how mutations affect transmissibility, antibody escape, lethality, and infectivity of these SARS-CoV-2 VOCs. By performing simulations of the trimeric ectodomains of the Spike^B.1.617.2^, Spike^BA.2^, Spike^BA.5^, Spike^BQ.1^ for 300 ns, we investigated the relationships of structural features of mutants and described these biological outcomes. D614G is related with the up- and down-RBD state conformations and P681H strongly correlates with low lethality. P681R, on the other hand, is associated with higher lethality and increased fusogenicity. Taken together, our results support the hypothesis that cooperativity mediated by residues D614G, P681R, and P681H impact both infectivity and lethality of SARS-CoV-2.

Phylogenetic tree of Spike^WT^, Spike^B.1.617.2^, Spike^BA.2^, Spike^BA.5^ and, Spike^BQ.1^ show two groups, in which Spike^B.1.617.2^ is closer related with Spike^WT^ than Omicron sub-variants (**Figure S3**). We hypothesize that SARS-CoV-2 evolved initially to increase interaction with hACE2 to increase the infection rate (Spike^B.1.617.2^) but the Omicron sub-variants seem to be involved to evade the immune system. We observed that the cooperativity between the three protomers within the trimeric Spike structure is lost mainly in the Omicron sub-variants. This observation may be important to turn off the communication between spike protomers in the infection process. Conversely, the Spike^Omicron^ mutations gain in the evasion of the immune system by hidden important epitopes. These observations are in agreement with previous studies, in which one-RBD-up conformation is maintained independent of hACE2 interaction probably due to the loss of protomer-protomer cooperativity. Furthermore, the immune system evasion is also increased in these Omicron sub-variants when compared to other VOCs (Z. Zhao et al., 2022). In addition, the observed mutations in Spike^Omicron^ analyzed in this study seem to favor the down-RBD state as a way of evading the immune system. At the same time, according to published studies such mutations do not significantly affect the interaction with hACE2 (Mannar et al., 2022). This supports the hypothesis that there is a new mechanism for activating the up-RBD state independent of cooperativity.

Previous study showed that the sub-variants BQ.1, BQ.1.1, BA.4.6, BF.7, and BA.2.75.2 are resistant to neutralizing antibodies from triple vaccinated individuals (Qu et al., 2022). However, all Omicron sub-variants maintained their weakened infectivity in the human lung cancer cell line (Calu-03), most notably the BA.2.75.2 sub-variant (Qu et al., 2022). The computational molecular dynamics presented here revealed significant structural changes in the S1 subunit in Spike^B.1.617.2^, Spike^BA.2^, Spike^BA.5^, and Spike^BQ.1^ (mainly in NTD and RBD). We also hypothesize that substitutions K417N, L452R, N444T, and N460K are able to promote resistance to neutralizing antibodies by improving their networks of hydrogen bonding interactions with neighboring residues, blocking interactions with neutralizing antibodies. These structural changes provide a basis for explaining the sensibility loss of neutralizing antibodies in VOCs. Our results also suggest mechanisms for how some mutations may contribute to evasion of the immune system in Delta and Omicron variants and may, therefore, help the study of updated vaccines against novel Omicron sub-lineages.

## Acknowledgment

The authors acknowledge the National Council for Scientific and Technological Development (CNPq), the Coordination for the Improvement of Higher Education Personnel (CAPES, grant 88887.374931/2019-00, Coordenação de Aperfeiçoamento de Pessoal de Nível Superior - Finance Code 01), and the São Paulo Research Foundation (FAPESP, grants 2019/00195-2, 2020/04680-0, 2016/09047-8), Rede Virus MCTI (grant FINEP 0459/20), Brazil, for financial support. The authors also acknowledge Geoambiente Sensoriamento Remoto LTDA for their generous support and for providing access to the Google Cloud services supported by grant FAPESP 2021/00070-5.

