## Supporting Information for "Cooperative and structural relationships of the trimeric Spike with infectivity and antibody escape of the strains Delta (B.1.617.2) and Omicron (BA.2, BA.5, and BQ.1)"

**Running title: Structural impacts of the different Spikes**

To whom correspondence should be addressed:

Cristiane R. Guzzo, Ph.D, Department of Microbiology, Institute of Biomedical Sciences, University of São Paulo, Av. Prof. Lineu Prestes, 1374, Cidade Universitária, 5508-900, São Paulo/SP, Brazil, +55 11 3091-7298;

Anacleto Silva de Souza, Ph.D, Department of Microbiology, Institute of Biomedical Sciences, University of São Paulo, Av. Prof. Lineu Prestes, 1374, Cidade Universitária, 5508-900, São Paulo/SP, Brazil, +55 11 3091-7298;

**Keywords:** B.1.617.2, BA.2, BA.5 and BQ.1; D614G; P681R; P681H; infectivity, transmissibility and immune system escape.

>WT
MFVFLVLLPLVSSQCVNLTTRTQLPPAYTNSFTRGVYYPDKVFRSSVLHSTQDLFLPFFSNVTWFHAIHVSGTNGTKRFDN
PVLPFNDGVYFASTEKSNIIRGWIFGTTLDSKTQSLLIVNNATNVVIKVCEFQFCNDPFLGVYYHKNNKSWMESEFRVYSS
ANNCTFEYVSQPFLMDLEGKQGNFKNLREFVFKNIDGYFKIYSKHTPINLVRDLPQGFSALEPLVDLPIGINITRFQTLLA
LHRSYLTPGDSSSGWTAGAAAYYVGYLQPRTFLLKYNENGTITDAVDCALDPLSETKCTLKSFTVEKGIYQTSNFRVQPTE
SIVRFPNITNLCPFGEVFNATRFASVYAWNRKRISNCVADYSVLYNSASFSTFKCYGVSPTKLNDLCFTNVYADSFVIRGD
EVRQIAPGQTGKIADYNYKLPDDFTGCVIAWNSNNLDSKVGGNYNYLYRLFRKSNLKPFERDISTEIYQAGSTPCNGVEGF
NCYFPLQSYGFQPTNGVGYQPYRVVVLSFELLHAPATVCGPKKSTNLVKNKCVNFNFNGLTGTGVLTESNKKFLPFQQFGR
DIADTTDAVRDPQTLEILDITPCSFGGVSVITPGTNTSNQVAVLYQDVNCTEVPVAIHADQLTPTWRVYSTGSNVFQTRAG
CLIGAEHVNNSYECDIPIGAGICASYQTQTNSPRRARSVASQSIIAYTMSLGAENSVAYSNNSIAIPTNFTISVTTEILPV
SMTKTSVDCTMYICGDSTECSNLLLQYGSFCTQLNRALTGIAVEQDKNTQEVFAQVKQIYKTPPIKDFGGFNFSQILPDPS
KPSKRSFIEDLLFNKVTLADAGFIKQYGDCLGDIAARDLICAQKFNGLTVLPPLLTDEMIAQYTSALLAGTITSGWTFGAG
AALQIPFAMQMAYRFNGIGVTQNVLYENQKLIANQFNSAIGKIQDSLSSTASALGKLQDVVNQNAQALNTLVKQLSSNFGA
ISSVLNDILSRLDKVEAEVQIDRLITGRLQSLQTYVTQQLIRAAEIRASANLAATKMSECVLGQSKRVDFCGKGYHLMSFP
QSAPHGVVFLHVTYVPAQEKNFTTAPAICHDGKAHFPREGVFVSNGTHWFVTQRNFYEPQIITTDNTFVSGNCDVVIGIVN
NTVYDPLQPELDSFKEELDKYFKNHTSPDVDLGDISGINASVVNIQKEIDRLNEVAKNLNESLIDLQELGKYEQYIKWPWY
IWLGFIAGLIAIVMVTIMLCCMTSCCSCLKGCCSCGSCCKFDEDDSEPVLKGVKLHYT

>B.1.617.2_EPI_ISL_15728122
MFVFLVLLPLVSSQCVNLRTRTQLPPAYTNSFTRGVYYPDKVFRSSVLHSTQDLFLPFFSNVTWFHAIHVSGTNRTKRFYN
PVLPFNDGVYFASIEKSNIIRGWIFGTTLDSKTQSLLIVNNATNVVIKVCEFQFCNDPFLDVYYHKNNKSWMESEFRVYSS
ANNCTFEYVSQPFLMDLEGKQGNFKNLREFVFKNIDGYFKIYSKHTPINLVRDLPQGFSALEPLVDLPIGINITRFQTLLA
LHRSYLTPGDSSSGWTAGAAAYYVGYLQPRTFLLKYNENGTITDAVDCALDPLSETKCTLKSFTVEKGIYQTSNFRVQPTE
SIVRFPNITNLCPFGEVFNATRFASVYAWNRKRISNCVADYSVLYNSASFSTFKCYGVSPTKLNDLCFTNVYADSFVIRGD
EVRQIAPGQTGKIADYNYKLPDDFTGCVIAWNSNNLDSKVGGNYNYRYRLFRKSNLKPFERDISTEIYQAGSKPCNGVEGF
NCYFPLQSYGFQPTNGVGYQPYRVVVLSFELLHAPATVCGPKKSTNLVKNKCVNFNFNGLTGTGVLTESNKKFLPFQQFGR
DIADTTDAVRDPQTLEILDITPCSFGGVSVITPGTNTSNQVAVLYQGVNCTEVPVAIHADQLTPTWRVYSTGSNVFQTRAG
CLIGAEHVNNSYECDIPIGAGICASYQTQTNSRRRARSVASQSIIAYTMSLGAENSVAYSNNSIAIPTNFTISVTTEILPV
SMTKTSVDCTMYICGDSTECSNLLLQYGSFCTQLNRALTGIAVEQDKNTQEVFAQVKQIYKTPPIKDFGGFNFSQILPDPS
KPSKRSFIEDLLFNKVTLADAGFIKQYGDCLGDIAARDLICAQKFNGLTVLPPLLTDEMIAQYTSALLAGTITSGWTFGAG
AALQIPFAMQMAYRFNGIGVTQNVLYENQKLIANQFNSAIGKIQDSLSSTASALGKLQNVVNQNAQALNTLVKQLSSNFGA
ISSVLNDILSRLDKVEAEVQIDRLITGRLQSLQTYVTQQLIRAAEIRASANLAATKMSECVLGQSKRVDFCGKGYHLMSFP
QSAPHGVVFLHVTYVPAQEKNFTTAPAICHDGKAHFPREGVFVSNGTHWFVTQRNFYEPQIITTDNTFVSGNCDVVIGIVN
NTVYDPLQPELDSFKEELDKYFKNHTSPDVDLGDISGINASVVNIQKEIDRLNEVAKNLNESLIDLQELGKYEQYIKWPWY
IWLGFIAGLIAIVMVTIMLCCMTSCCSCLKGCCSCGSCCKFDEDDSEPLLKGVKLHYT

>BA.2_EPI_ISL_12905607
MFVFLVLLPLVSSQCVNLITRTQSYTNSFTRGVYYPDKVFRSSVLHSTQDLFLPFFSNVTWFHAIHVSGTNGTKRFDNPVL
PFNDGVYFASTEKSNIIRGWIFGTTLDSKTQSLLIVNNATNVVIKVCEFQFCNDPFLDVYYHKNNKSWMESEFRVYSSANN
CTFEYVSQPFLMDLEGKQGNFKNLREFVFKNIDGYFKIYSKHTPINLGRDLPQGFSALEPLVDLPIGINITRFQTLLALHR
SYLTPGDSSSGWTAGAAAYYVGYLQPRTFLLKYNENGTITDAVDCALDPLSETKCTLKSFTVEKGIYQTSNFRVQPTESIV
RFPNITNLCPFDEVFNATRFASVYAWNRKRISNCVADYSVLYNFAPFFAFKCYGVSPTKLNDLCFTNVYADSFVIRGNEVS
QIAPGQTGNIADYNYKLPDDFTGCVIAWNSNNLDSKVGGNYNYLYRLFRKSNLKPFERDISTEIYQAGNKPCNGVAGFNCY
FPLRSYGFRPTYGVGHQPYRVVVLSFELLHAPATVCGPKKSTNLVKNKCVNFNFNGLTGTGVLTESNKKFLPFQQFGRDIA
DTTDAVRDPQTLEILDITPCSFGGVSVITPGTNTSNQVAVLYQGVNCTEVPVAIHADQLTPTWRVYSTGSNVFQTRAGCLI
GAEYVNNSYECDIPIGAGICASYQTQTKSHRRARSVASQSIIAYTMSLGAENSVAYSNNSIAIPTNFTISVTTEILPVSMT
KTSVDCTMYICGDSTECSNLLLQYGSFCTQLKRALTGIAVEQDKNTQEVFAQVKQIYKTPPIKYFGGFNFSQILPDPSKPS
KRSFIEDLLFNKVTLADAGFIKQYGDCLGDIAARDLICAQKFNGLTVLPPLLTDEMIAQYTSALLAGTITSGWTFGAGAAL
QIPFAMQMAYRFNGIGVTQNVLYENQKLIANQFNSAIGKIQDSLSSTASALGKLQDVVNHNAQALNTLVKQLSSKFGAISS
VLNDILSRLDKVEAEVQIDRLITGRLQSLQTYVTQQLIRAAEIRASANLAATKMSECVLGQSKRVDFCGKGYHLMSFPQSA
PHGVVFLHVTYVPAQEKNFTTAPAICHDGKAHFPREGVFVSNGTHWFVTQRNFYEPQIITTDNTFVSGNCDVVIGIVNNTV
YDPLQPELDSFKEELDKYFKNHTSPDVDLGDISGINASVVNIQKEIDRLNEVAKNLNESLIDLQELGKYEQYIKWPWYIWL
GFIAGLIAIVMVTIMLCCMTSCCSCLKGCCSCGSCCKFDEDDSEPVLKGVKLHYT

>BA.5_EPI_ISL_14439279
MFVFLVLLPLVSSQCVNLITRTQSYTNSFTRGVYYPDKVFRSSVLHSTQDLFLPFFSNVTWFHAISGTNGTKRFDNPVLPF
NDGVYFASTEKSNIIRGWIFGTTLDSKTQSLLIVNNATNVVIKVCEFQFCNDPFLDVYYHKNNKSWMESEFRVYSSANNCT
FEYVSQPFLMDLEGKQGNFKNLREFVFKNIDGYFKIYSKHTPINLGRDLPQGFSALEPLVDLPIGINITRFQTLLALHRSY
LTPGDSSSGWTAGAAAYYVGYLQPRTFLLKYNENGTITDAVDCALDPLSETKCTLKSFTVEKGIYQTSNFRVQPTESIVRF
PNITNLCPFDEVFNATRFASVYAWNRKRISNCVADYSVLYNFAPFFAFKCYGVSPTKLNDLCFTNVYADSFVIRGNEVSQI

```
APGQTGNIADYNYKLPDDFTGCVIAWNSNKLDSKVGGNYNYRYRLFRKSNLKPFERDISTEIYQAGNKPCNGVAGVNCYFP
LQSYGFRPTYGVGHQPYRVVVLSFELLHAPATVCGPKKSTNLVKNKCVNFNFNGLTGTGVLTESNKKFLPFQQFGRDIADT
TDAVRDPQTLEILDITPCSFGGVSVITPGTNTSNQVAVLYQGVNCTEVPVAIHADQLTPTWRVYSTGSNVFQTRAGCLIGA
EYVNNSYECDIPIGAGICASYQTQTKSHRRARSVASQSIIAYTMSLGAENSVAYSNNSIAIPTNFTISVTTEILPVSMTKT
SVDCTMYICGDSTECSNLLLQYGSFCTQLKRALTGIAVEQDKNTQEVFAQVKQIYKTPPIKYFGGFNFSQILPDPSKPSKR
SFIEDLLFNKVTLADAGFIKQYGDCLGDIAARDLICAQKFNGLTVLPPLLTDEMIAQYTSALLAGTITSGWTFGAGAALQI
PFAMQMAYRFNGIGVTQNVLYENQKLIANQFNSAIGKIQDSLSSTASALGKLQDVVNHNAQALNTLVKQLSSKFGAISSVL
NDILSRLDKVEAEVQIDRLITGRLQSLQTYVTQQLIRAAEIRASANLAATKMSECVLGQSKRVDFCGKGYHLMSFPQSAPH
GVVFLHVTYVPAQEKNFTTAPAICHDGKAHFPREGVFVSNGTHWFVTQRNFYEPQIITTDNTFVSGNCDVVIGIVNNTVYD
PLQPELDSFKEELDKYFKNHTSPDVDLGDISGINASVVNIQKEIDRLNEVAKNLNESLIDLQELGKYEQYIKWPWYIWLGF
IAGLIAIVMVTIMLCCMTSCCSCLKGCCSCGSCCKFDEDDSEPVLKGVKLHYT

>BQ.1_EPI_ISL_15731012
MFVFLVLLPLVSSQCVNLITRTQSYTNSFTRGVYYPDKVFRSSVLHSTQDLFLPFFSNVTWFHAISGTNGTKRFDNPVLPF
NDGVYFASTEKSNIIRGWIFGTTLDSKTQSLLIVNNATNVVIKVCEFQFCNDPFLDVYHKNNKSWMESEFRVYSSANNCTF
EYVSQPFLMDLEGKQGNFKNLREFVFKNIDGYFKIYSKHTPINLGRDLPQGFSALEPLVDLPIGINITRFQTLLALHRSYL
TPGDSSSGWTAGAAAYYVGYLQPRTFLLKYNENGTITDAVDCALDPLSETKCTLKSFTVEKGIYQTSNFRVQPTESIVRFP
NITNLCPFDEVFNATRFASVYAWNRKRISNCVADYSVLYNFAPFFAFKCYGVSPTKLNDLCFTNVYADSFVIRGNEVSQIA
PGQTGNIADYNYKLPDDFTGCVIAWNSNKLDSTVGGNYNYRYRLFRKSKLKPFERDISTEIYQAGNKPCNGVAGVNCYFPL
QSYGFRPTYGVGHQPYRVVVLSFELLHAPATVCGPKKSTNLVKNKCVNFNFNGLTGTGVLTESNKKFLPFQQFGRDIADTT
DAVRDPQTLEILDITPCSFGGVSVITPGTNTSNQVAVLYQGVNCTEVPVAIHADQLTPTWRVYSTGSNVFQTRAGCLIGAE
YVNNSYECDIPIGAGICASYQTQTKSHRRARSVASQSIIAYTMSLGAENSVAYSNNSIAIPTNFTISVTTEILPVSMTKTS
VDCTMYICGDSTECSNLLLQYGSFCTQLKRALTGIAVEQDKNTQEVFAQVKQIYKTPPIKYFGGFNFSQILPDPSKPSKRS
FIEDLLFNKVTLADAGFIKQYGDCLGDIAARDLICAQKFNGLTVLPPLLTDEMIAQYTSALLAGTITSGWTFGAGAALQIP
FAMQMAYRFNGIGVTQNVLYENQKLIANQFNSAIGKIQDSLSSTASALGKLQDVVNHNAQALNTLVKQLSSKFGAISSVLN
DILSRLDKVEAEVQIDRLITGRLQSLQTYVTQQLIRAAEIRASANLAATKMSECVLGQSKRVDFCGKGYHLMSFPQSAPHG
VVFLHVTYVPAQEKNFTTAPAICHDGKAHFPREGVFVSNGTHWFVTQRNFYEPQIITTDNTFVSGNCDVVIGIVNNTVYDP
LQPELDSFKEELDKYFKNHTSPDVDLGDISGINASVVNIQKEIDRLNEVAKNLNESLIDLQELGKYEQYIKWPWYIWLGFI
AGLIAIVMVTIMLCCMTSCCSCLKGCCSCGSCCKFDEDDSEPVLKGVKLHYT
```

**Figure S1.** Primary sequences of Spike protein obtained from Translate Tools available in Expasy web-service (WT, BA.2, BA.5 and BQ.1). The genomic sequences of these sub-lineage were obtained from GISAID initiative using the accession code EPI_ISL_402123.1, EPI_ISL_15728122, EPI_ISL_12905607, EPI_ISL_14439279 and EPI_ISL_15731012 and translated in amino acids using Translate Tool website (ExPASy, 2022).

```
                                            14     19       27
                                             |     |        |
WT                MFLLTTKRTMFVFLVLLPLVSSQCVNLTTRTQLPPAYTNSFTRGVYYPDKVFRSSVLHST
B.1.617.2         MFLLTTKRTMFVFLVLLPLVSSQCVNLRTRTQLPPAYTNSFTRGVYYPDKVFRSSVLHST
BA.2              MFLLTTKRTMFVFLVLLPLVSSQCVNLITRTQ---SYTNSFTRGVYYPDKVFRSSVLHST
BA.5              MFLLTTKRTMFVFLVLLPLVSSQCVNLITRTQ---SYTNSFTRGVYYPDKVFRSSVLHST
BQ.1              MFLLTTKRTMFVFLVLLPLVSSQCVNLITRTQ---SYTNSFTRGVYYPDKVFRSSVLHST
                  *************************** ****   :************************

                                       69     75    80                95
                                       |      |     |                 |
WT                QDLFLPFFSNVTWFHAIHVSGTNGTKRFDNPVLPFNDGVYFASTEKSNIIRGWIFGTTLD
B.1.617.2         QDLFLPFFSNVTWFHAIHVSGTNRTKRFYNPVLPFNDGVYFASIEKSNIIRGWIFGTTLD
BA.2              QDLFLPFFSNVTWFHAIHVSGTNGTKRFDNPVLPFNDGVYFASTEKSNIIRGWIFGTTLD
BA.5              QDLFLPFFSNVTWFHAI--SGTNGTKRFDNPVLPFNDGVYFASTEKSNIIRGWIFGTTLD
BQ.1              QDLFLPFFSNVTWFHAI--SGTNGTKRFDNPVLPFNDGVYFASTEKSNIIRGWIFGTTLD
                  *****************  **** **** ************** ****************
```

```
                                             142
                                              |
WT              SKTQSLLIVNNATNVVIKVCEFQFCNDPFLGVYYHKNNKSWMESEFRVYSSANNCTFEYV
B.1.617.2       SKTQSLLIVNNATNVVIKVCEFQFCNDPFLDVYYHKNNKSWMESEFRVYSSANNCTFEYV
BA.2            SKTQSLLIVNNATNVVIKVCEFQFCNDPFLDVYYHKNNKSWMESEFRVYSSANNCTFEYV
BA.5            SKTQSLLIVNNATNVVIKVCEFQFCNDPFLDVYYHKNNKSWMESEFRVYSSANNCTFEYV
BQ.1            SKTQSLLIVNNATNVVIKVCEFQFCNDPFLDVY-HKNNKSWMESEFRVYSSANNCTFEYV
                ******************************.** **************************

                                                        213
                                                         |
WT              SQPFLMDLEGKQGNFKNLREFVFKNIDGYFKIYSKHTPINLVRDLPQGFSALEPLVDLPI
B.1.617.2       SQPFLMDLEGKQGNFKNLREFVFKNIDGYFKIYSKHTPINLVRDLPQGFSALEPLVDLPI
BA.2            SQPFLMDLEGKQGNFKNLREFVFKNIDGYFKIYSKHTPINLGRDLPQGFSALEPLVDLPI
BA.5            SQPFLMDLEGKQGNFKNLREFVFKNIDGYFKIYSKHTPINLGRDLPQGFSALEPLVDLPI
BQ.1            SQPFLMDLEGKQGNFKNLREFVFKNIDGYFKIYSKHTPINLGRDLPQGFSALEPLVDLPI
                ***************************************** ******************

WT              GINITRFQTLLALHRSYLTPGDSSSGWTAGAAAYYVGYLQPRTFLLKYNENGTITDAVDC
B.1.617.2       GINITRFQTLLALHRSYLTPGDSSSGWTAGAAAYYVGYLQPRTFLLKYNENGTITDAVDC
BA.2            GINITRFQTLLALHRSYLTPGDSSSGWTAGAAAYYVGYLQPRTFLLKYNENGTITDAVDC
BA.5            GINITRFQTLLALHRSYLTPGDSSSGWTAGAAAYYVGYLQPRTFLLKYNENGTITDAVDC
BQ.1            GINITRFQTLLALHRSYLTPGDSSSGWTAGAAAYYVGYLQPRTFLLKYNENGTITDAVDC
                ************************************************************

                       300                 320                339
                        |                   |                  |
WT              ALDPLSETKCTLKSFTVEKGIYQTSNFRVQPTESIVRFPNITNLCPFGEVFNATRFASVY
B.1.617.2       ALDPLSETKCTLKSFTVEKGIYQTSNFRVQPTESIVRFPNITNLCPFGEVFNATRFASVY
BA.2            ALDPLSETKCTLKSFTVEKGIYQTSNFRVQPTESIVRFPNITNLCPFDEVFNATRFASVY
BA.5            ALDPLSETKCTLKSFTVEKGIYQTSNFRVQPTESIVRFPNITNLCPFDEVFNATRFASVY
BQ.1            ALDPLSETKCTLKSFTVEKGIYQTSNFRVQPTESIVRFPNITNLCPFDEVFNATRFASVY
                ***********************************************.************

                                  371                               405
                                   |                                 |
WT              AWNRKRISNCVADYSVLYNSASFSTFKCYGVSPTKLNDLCFTNVYADSFVIRGDEVRQIA
B.1.617.2       AWNRKRISNCVADYSVLYNSASFSTFKCYGVSPTKLNDLCFTNVYADSFVIRGDEVRQIA
BA.2            AWNRKRISNCVADYSVLYNFAPFFAFKCYGVSPTKLNDLCFTNVYADSFVIRGNEVSQIA
BA.5            AWNRKRISNCVADYSVLYNFAPFFAFKCYGVSPTKLNDLCFTNVYADSFVIRGNEVSQIA
BQ.1            AWNRKRISNCVADYSVLYNFAPFFAFKCYGVSPTKLNDLCFTNVYADSFVIRGNEVSQIA
                ******************* * * :****************************:** ***

                    417                    440 444     452     460
                     |                      |   |       |       |
WT              PGQTGKIADYNYKLPDDFTGCVIAWNSNNLDSKVGGNYNYLYRLFRKSNLKPFERDISTE
B.1.617.2       PGQTGKIADYNYKLPDDFTGCVIAWNSNNLDSKVGGNYNYRYRLFRKSNLKPFERDISTE
BA.2            PGQTGNIADYNYKLPDDFTGCVIAWNSNNLDSKVGGNYNYLYRLFRKSNLKPFERDISTE
BA.5            PGQTGNIADYNYKLPDDFTGCVIAWNSNKLDSKVGGNYNYRYRLFRKSNLKPFERDISTE
BQ.1            PGQTGNIADYNYKLPDDFTGCVIAWNSNKLDSTVGGNYNYRYRLFRKSKLKPFERDISTE
                *****:**********************:***.******* *******:***********

                    477    484      493  498    505
                     |      |        |    |      |
WT              IYQAGSTPCNGVEGFNCYFPLQSYGFQPTNGVGYQPYRVVVLSFELLHAPATVCGPKKST
B.1.617.2       IYQAGSKPCNGVEGFNCYFPLQSYGFQPTNGVGYQPYRVVVLSFELLHAPATVCGPKKST
BA.2            IYQAGNKPCNGVAGFNCYFPLRSYGFRPTYGVGHQPYRVVVLSFELLHAPATVCGPKKST
BA.5            IYQAGNKPCNGVAGVNCYFPLQSYGFRPTYGVGHQPYRVVVLSFELLHAPATVCGPKKST
BQ.1            IYQAGNKPCNGVAGVNCYFPLQSYGFRPTYGVGHQPYRVVVLSFELLHAPATVCGPKKST
                *****..***** *.******:****:** ***:**************************

WT              NLVKNKCVNFNFNGLTGTGVLTESNKKFLPFQQFGRDIADTTDAVRDPQTLEILDITPCS
B.1.617.2       NLVKNKCVNFNFNGLTGTGVLTESNKKFLPFQQFGRDIADTTDAVRDPQTLEILDITPCS
BA.2            NLVKNKCVNFNFNGLTGTGVLTESNKKFLPFQQFGRDIADTTDAVRDPQTLEILDITPCS
BA.5            NLVKNKCVNFNFNGLTGTGVLTESNKKFLPFQQFGRDIADTTDAVRDPQTLEILDITPCS
BQ.1            NLVKNKCVNFNFNGLTGTGVLTESNKKFLPFQQFGRDIADTTDAVRDPQTLEILDITPCS
```

```
               ************************************************************

                                     614
                                     |
WT             FGGVSVITPGTNTSNQVAVLYQDVNCTEVPVAIHADQLTPTWRVYSTGSNVFQTRAGCLI
B.1.617.2      FGGVSVITPGTNTSNQVAVLYQGVNCTEVPVAIHADQLTPTWRVYSTGSNVFQTRAGCLI
BA.2           FGGVSVITPGTNTSNQVAVLYQGVNCTEVPVAIHADQLTPTWRVYSTGSNVFQTRAGCLI
BA.5           FGGVSVITPGTNTSNQVAVLYQGVNCTEVPVAIHADQLTPTWRVYSTGSNVFQTRAGCLI
BQ.1           FGGVSVITPGTNTSNQVAVLYQGVNCTEVPVAIHADQLTPTWRVYSTGSNVFQTRAGCLI
               **********************.*************************************

                  655                     679
                  |                       |
WT             GAEHVNNSYECDIPIGAGICASYQTQTNSPRRARSVASQSIIAYTMSLGAENSVAYSNNS
B.1.617.2      GAEHVNNSYECDIPIGAGICASYQTQTNSRRRARSVASQSIIAYTMSLGAENSVAYSNNS
BA.2           GAEYVNNSYECDIPIGAGICASYQTQTKSHRRARSVASQSIIAYTMSLGAENSVAYSNNS
BA.5           GAEYVNNSYECDIPIGAGICASYQTQTKSHRRARSVASQSIIAYTMSLGAENSVAYSNNS
BQ.1           GAEYVNNSYECDIPIGAGICASYQTQTKSHRRARSVASQSIIAYTMSLGAENSVAYSNNS
               ***:***********************:* ******************************

                                                                   764
                                                                   |
WT             IAIPTNFTISVTTEILPVSMTKTSVDCTMYICGDSTECSNLLLQYGSFCTQLNRALTGIA
B.1.617.2      IAIPTNFTISVTTEILPVSMTKTSVDCTMYICGDSTECSNLLLQYGSFCTQLNRALTGIA
BA.2           IAIPTNFTISVTTEILPVSMTKTSVDCTMYICGDSTECSNLLLQYGSFCTQLKRALTGIA
BA.5           IAIPTNFTISVTTEILPVSMTKTSVDCTMYICGDSTECSNLLLQYGSFCTQLKRALTGIA
BQ.1           IAIPTNFTISVTTEILPVSMTKTSVDCTMYICGDSTECSNLLLQYGSFCTQLKRALTGIA
               ****************************************************:*******

                                       796                815
                                       |                  |
WT             VEQDKNTQEVFAQVKQIYKTPPIKDFGGFNFSQILPDPSKPSKRSFIEDLLFNKVTLADA
B.1.617.2      VEQDKNTQEVFAQVKQIYKTPPIKDFGGFNFSQILPDPSKPSKRSFIEDLLFNKVTLADA
BA.2           VEQDKNTQEVFAQVKQIYKTPPIKYFGGFNFSQILPDPSKPSKRSFIEDLLFNKVTLADA
BA.5           VEQDKNTQEVFAQVKQIYKTPPIKYFGGFNFSQILPDPSKPSKRSFIEDLLFNKVTLADA
BQ.1           VEQDKNTQEVFAQVKQIYKTPPIKYFGGFNFSQILPDPSKPSKRSFIEDLLFNKVTLADA
               ************************ ***********************************

WT             GFIKQYGDCLGDIAARDLICAQKFNGLTVLPPLLTDEMIAQYTSALLAGTITSGWTFGAG
B.1.617.2      GFIKQYGDCLGDIAARDLICAQKFNGLTVLPPLLTDEMIAQYTSALLAGTITSGWTFGAG
BA.2           GFIKQYGDCLGDIAARDLICAQKFNGLTVLPPLLTDEMIAQYTSALLAGTITSGWTFGAG
BA.5           GFIKQYGDCLGDIAARDLICAQKFNGLTVLPPLLTDEMIAQYTSALLAGTITSGWTFGAG
BQ.1           GFIKQYGDCLGDIAARDLICAQKFNGLTVLPPLLTDEMIAQYTSALLAGTITSGWTFGAG
               ************************************************************

WT             AALQIPFAMQMAYRFNGIGVTQNVLYENQKLIANQFNSAIGKIQDSLSSTASALGKLQDV
B.1.617.2      AALQIPFAMQMAYRFNGIGVTQNVLYENQKLIANQFNSAIGKIQDSLSSTASALGKLQNV
BA.2           AALQIPFAMQMAYRFNGIGVTQNVLYENQKLIANQFNSAIGKIQDSLSSTASALGKLQDV
BA.5           AALQIPFAMQMAYRFNGIGVTQNVLYENQKLIANQFNSAIGKIQDSLSSTASALGKLQDV
BQ.1           AALQIPFAMQMAYRFNGIGVTQNVLYENQKLIANQFNSAIGKIQDSLSSTASALGKLQDV
               **********************************************************:*

                 954            969              986           1000
                 |              |                |             |
WT             VNQNAQALNTLVKQLSSNFGAISSVLNDILSRLDKVEAEVQIDRLITGRLQSLQTYVTQQ
B.1.617.2      VNQNAQALNTLVKQLSSNFGAISSVLNDILSRLDKVEAEVQIDRLITGRLQSLQTYVTQQ
BA.2           VNHNAQALNTLVKQLSSKFGAISSVLNDILSRLDKVEAEVQIDRLITGRLQSLQTYVTQQ
BA.5           VNHNAQALNTLVKQLSSKFGAISSVLNDILSRLDKVEAEVQIDRLITGRLQSLQTYVTQQ
BQ.1           VNHNAQALNTLVKQLSSKFGAISSVLNDILSRLDKVEAEVQIDRLITGRLQSLQTYVTQQ
               **:**************:******************************************

WT             LIRAAEIRASANLAATKMSECVLGQSKRVDFCGKGYHLMSFPQSAPHGVVFLHVTYVPAQ
B.1.617.2      LIRAAEIRASANLAATKMSECVLGQSKRVDFCGKGYHLMSFPQSAPHGVVFLHVTYVPAQ
BA.2           LIRAAEIRASANLAATKMSECVLGQSKRVDFCGKGYHLMSFPQSAPHGVVFLHVTYVPAQ
BA.5           LIRAAEIRASANLAATKMSECVLGQSKRVDFCGKGYHLMSFPQSAPHGVVFLHVTYVPAQ
```

```
BQ.1            LIRAAEIRASANLAATKMSECVLGQSKRVDFCGKGYHLMSFPQSAPHGVVFLHVTYVPAQ
                ************************************************************

WT              EKNFTTAPAICHDGKAHFPREGVFVSNGTHWFVTQRNFYEPQIITTDNTFVSGNCDVVIG
B.1.617.2       EKNFTTAPAICHDGKAHFPREGVFVSNGTHWFVTQRNFYEPQIITTDNTFVSGNCDVVIG
BA.2            EKNFTTAPAICHDGKAHFPREGVFVSNGTHWFVTQRNFYEPQIITTDNTFVSGNCDVVIG
BA.5            EKNFTTAPAICHDGKAHFPREGVFVSNGTHWFVTQRNFYEPQIITTDNTFVSGNCDVVIG
BQ.1            EKNFTTAPAICHDGKAHFPREGVFVSNGTHWFVTQRNFYEPQIITTDNTFVSGNCDVVIG
                ************************************************************

WT              IVNNTVYDPLQPELDSFKEELDKYFKNHTSPDVDLGDISGINASVVNIQKEIDRLNEVAK
B.1.617.2       IVNNTVYDPLQPELDSFKEELDKYFKNHTSPDVDLGDISGINASVVNIQKEIDRLNEVAK
BA.2            IVNNTVYDPLQPELDSFKEELDKYFKNHTSPDVDLGDISGINASVVNIQKEIDRLNEVAK
BA.5            IVNNTVYDPLQPELDSFKEELDKYFKNHTSPDVDLGDISGINASVVNIQKEIDRLNEVAK
BQ.1            IVNNTVYDPLQPELDSFKEELDKYFKNHTSPDVDLGDISGINASVVNIQKEIDRLNEVAK
                ************************************************************


WT              NLNESLIDLQELGKYEQYIKWPWYIWLGFIAGLIAIVMVTIMLCCMTSCCSCLKGCCSCG
B.1.617.2       NLNESLIDLQELGKYEQYIKWPWYIWLGFIAGLIAIVMVTIMLCCMTSCCSCLKGCCSCG
BA.2            NLNESLIDLQELGKYEQYIKWPWYIWLGFIAGLIAIVMVTIMLCCMTSCCSCLKGCCSCG
BA.5            NLNESLIDLQELGKYEQYIKWPWYIWLGFIAGLIAIVMVTIMLCCMTSCCSCLKGCCSCG
BQ.1            NLNESLIDLQELGKYEQYIKWPWYIWLGFIAGLIAIVMVTIMLCCMTSCCSCLKGCCSCG
                ************************************************************


WT              SCCKFDEDDSEPVLKGVKLHYT
B.1.617.2       SCCKFDEDDSEPLLKGVKLHYT
BA.2            SCCKFDEDDSEPVLKGVKLHYT
BA.5            SCCKFDEDDSEPVLKGVKLHYT
BQ.1            SCCKFDEDDSEPVLKGVKLHYT
                ************:*********
```

**Figure S2. Multiple sequence alignment of Spike$^{WT}$, Spike$^{B.1.617.2}$, Spike$^{BA.2}$, Spike$^{BA.5}$ and Spike$^{BQ.1}$**. Amino acid sequence alignment of Spike$^{WT}$, Spike$^{B.1.617.2}$, Spike$^{BA.2}$, Spike$^{BA.5}$ and Spike$^{BQ.1}$ performed in the Multiple Alignment using Fast Fourier Transform (MAFF) (https://www.ebi.ac.uk/Tools/msa/mafft). Residues in bold represent the signal sequence (SS).

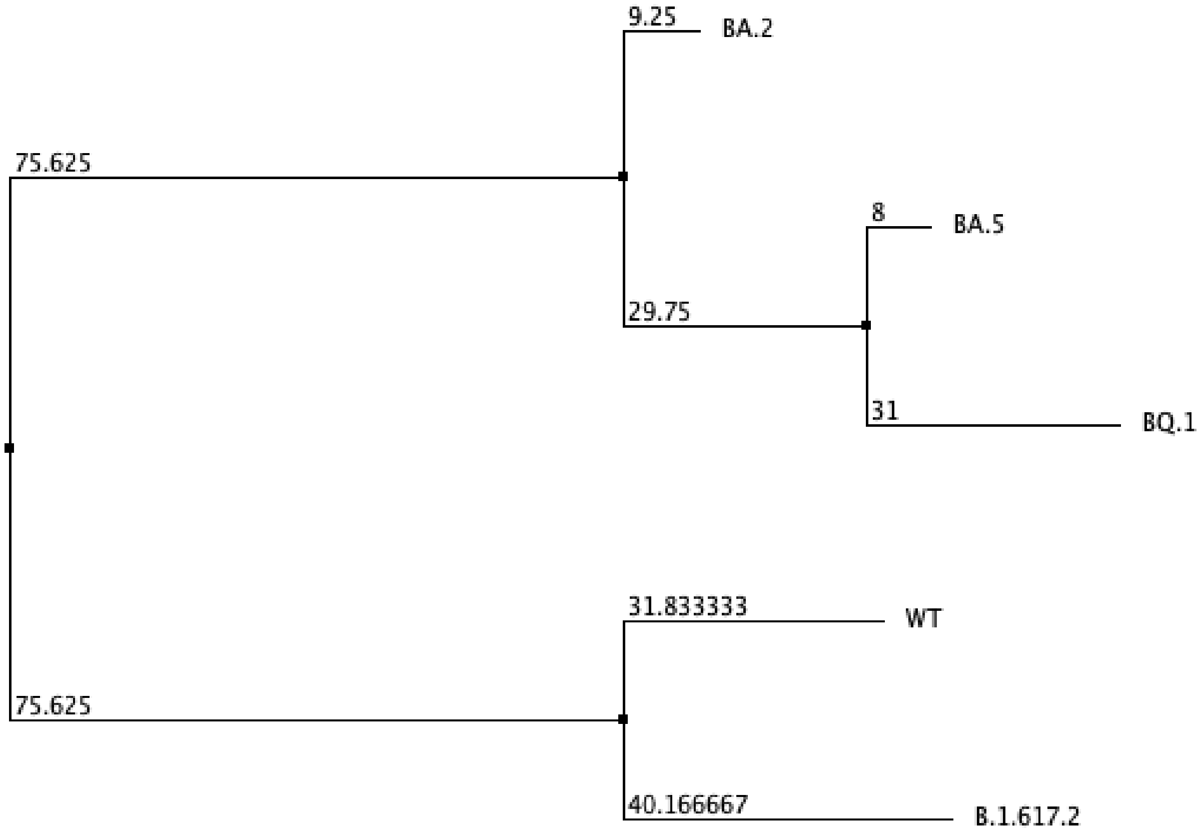


**Figure S3. Phylogenetic tree of Spike$^{WT}$, Spike$^{B.1.617.2}$, Spike$^{BA.2}$, Spike$^{BA.5}$ and Spike$^{BQ.1}$**. From a multiple sequence alignment was performed using the Jalview program, using Tcoffee followed by a phylogenetic tree calculation using Neighbour joining and Blosum62 matrix (Waterhouse et al., 2009).

**Table S1. Effect of K417N mutation in the hydrogen bonding networks.** Relevant hydrogen bonding interactions were identified in molecular dynamics, quantifying average hydrogen bonding occupancies from chains A, B and C of the residues K417 (Delta variant; B.1.617.2), N414 (BA.2), N412 (BA.5), and N411 (BQ.1) with neighboring residues.

| Residue | Hydrogen bonding occupancies |
|---|---|
| $_{SC}$K417$^{B.1.617.2}$ | $_{SC}$N422 (35.96%), $_{SC}$E406 (76.32%), $_{SC}$Y421 (27.40%), $_{MC}$F374 (30.24%) |
| $_{SC}$N414$^{BA.2}$ (K417N) | $_{SC}$N367$^{BA.2}$ (33.41%) and $_{MC}$R451$^{BA.2}$ (62.75%) |
| $_{SC}$N412$^{BA.5}$ (K417N) | $_{SC}$Q404$^{BA.5}$ (17.37%), $_{MC}$R449$^{BA.5}$ (66.12%), and $_{SC}$Y416$^{BA.5}$ (10.72%) |
| $_{SC}$N411$^{BQ.1}$ (K417N) | $_{SC}$E409$^{BQ.1}$ (28.68%), $_{MC}$A375$^{BQ.1}$ (15.03%), $_{MC}$Y372$^{BQ.1}$ (23.51%), $_{MC}$R457$^{BQ.1}$ (49.13%), $_{MC}$L458$^{BQ.1}$ (10.31%), and $_{SC}$Y424$^{BQ.1}$ (12.36%) |

Cut-off of distance and angle of 4 Å and 20°, respectively.

**Table S2. Effect of L452R mutation in the hydrogen bonding networks.** Relevant hydrogen bonding interactions were identified in molecular dynamics, quantifying average hydrogen bonding occupancies from chains A, B and C of the residues R452 (Delta variant; B.1.617.2), R447 (BA.5), and R445 (BQ.1) with neighboring residues.

| Residue | Hydrogen bonding occupancies |
|---|---|
| $_{SC}$R452$^{B.1.617.2}$ | $_{SC}$Y351$^{B.1.617.2}$ (18.41%), $_{MC}$N450$^{B.1.617.2}$ (23.53%), $_{SC}$E484$^{B.1.617.2}$ (87.43%), and $_{SC}$S494$^{SC}$ (9.43%) |
| $_{SC}$R447$^{BA.5}$ | $_{SC}$Y346$^{BA.5}$ (35.37%), $_{MC}$N445$^{BA.5}$ (29.54%), $_{SC}$S489$^{BA.5}$ (18.39%), and $_{MC}$S489$^{BA.5}$ (14.23%) |
| $_{SC}$R445$^{BQ.1}$ | $_{SC}$D436$^{BQ.1}$ (86.92%), $_{MC}$S437$^{BQ.1}$ (18.56%), $_{SC}$F341$^{BQ.1}$ (35.88%), $_{SC}$N444$^{BQ.1}$ (43.39%), $_{SC}$Y489$^{BQ.1}$ (20.50%), and $_{MC}$Y489$^{BQ.1}$ (16.10%) |

Cut-off of distance and angle of 4 Å and 20°, respectively. SC = side chain; MC = main chain.

**Table S3. Effect of K444T mutation in the hydrogen bonding networks.** Relevant hydrogen bonding interactions were identified in molecular dynamics, quantifying average hydrogen bonding occupancies from chains A, B and C of the residues K439 (BA.5), and T438 (BQ.1) with neighboring residues.

| Residue | Hydrogen bonding occupancies |
|---|---|
| $_{SC}$K439$^{BA.5}$ | $_{MC}$N443$^{BA.5}$ (11.29%), $_{MC}$G442$^{BA.5}$ (22.57%), and $_{SC}$N443$^{BA.5}$ (10.56%) |
| $_{SC}$T438$^{BQ.1}$ | $_{MC}$S432$^{BQ.1}$ (14.12%), $_{MC}$D436$^{BQ.1}$ (12.49%), $_{SC}$N442$^{BQ.1}$ (43.39%), and $_{SC}$R503$^{BQ.1}$ (53.53%) |

Cut-off of distance and angle of 4 Å and 20°, respectively. SC = side chain; MC = main chain.

**Table S4. Effect of N460K mutation in the hydrogen bonding networks.** Relevant hydrogen bonding interactions were identified in molecular dynamics, quantifying average hydrogen bonding occupancies from chains A, B and C of the residues N460 (Delta variant; B.1.617.2), N457 (BA.2), N455 (BA.5) and K454 (BQ.1) with neighboring residues.

| Residue | Hydrogen bonding occupancies |
|---|---|
| $_{SC}$N460$^{B.1.617.2}$ | $_{SC}$D420$^{B.1.617.2}$ (80.47%), $_{SC}$K424$^{B.1.617.2}$ (62.39%), and $_{SC}$Y421$^{B.1.617.2}$ (39.35%) |
| $_{SC}$N457$^{BA.2}$ | $_{C}$D417$^{BA.2}$ (74.96%), and $_{SC}$K421$^{BA.2}$ (63.93%) |
| $_{SC}$N455$^{BA.5}$ | $_{SC}$K419$^{BA.5}$ (38.71%), $_{SC}$D415$^{BA.5}$ (57.35%) and $_{SC}$Y416$^{BA.5}$ (10.11%) |
| $_{SC}$K454$^{BQ.1}$ | $_{SC}$D979$^{BQ.1}$ (35.07%), $_{SC}$D414$^{BQ.1}$ (94.89%) and $_{SC}$Y415$^{BQ.1}$ (38.01%) |

Cut-off of distance and angle of 4 Å and 20°, respectively. SC = side chain; MC = main chain.
